# How understudied populations have contributed to our understanding of Alzheimer’s disease genetics

**DOI:** 10.1101/2020.06.11.146993

**Authors:** Nadia Dehghani, Jose Bras, Rita Guerreiro

**Affiliations:** Center for Neurodegenerative Science, Van Andel Institute, Grand Rapids, Michigan, USA; Division of Psychiatry and Behavioral Medicine, Michigan State University College of Human Medicine, Grand Rapids, MI, USA

**Author notes:** Corresponding author Rita Guerreiro, Center for Neurodegenerative Science, Van Andel Institute, 333 Bostwick Ave. N.E., Grand Rapids, Michigan 49503-2518 USA.

**Keywords:** Alzheimer’s disease, genetics, understudied populations, diversity

## Abstract

The majority of genome-wide association studies have been conducted using samples with a European genetic background. As a field, we acknowledge this limitation and the need to increase the diversity of populations studied. A major challenge when designing and conducting such studies is to assimilate large samples sizes so that we attain enough statistical power to detect variants associated with disease, particularly when trying to identify variants with low and rare minor allele frequencies. In this study, we aimed to illustrate the benefits, to genetic characterization of Alzheimer’s disease (AD), in researching currently understudied populations. This is important for both fair representation of world populations and the translatability of findings. To that end, we have conducted a literature search to understand the contributions of studies, on different populations, to AD genetics. We systematically quantified the number of studies identifying mutations in known disease-causing genes, in a world-wide manner, and discussed the contributions of research in understudied populations to the identification of novel genetic factors in this disease. Additionally, we compared the effects of genome-wide significant SNPs across populations by focusing on loci that show different association profiles between populations (a key example being *APOE*). This work functions to both highlight how understudied populations have furthered our understanding of AD genetics, and to help us gage our progress in understanding the genetic architecture of this disease in all populations.

## Introduction

The purpose of this review is to highlight key findings on the genetics of Alzheimer’s disease (AD) from studies performed on understudied populations. We define such populations in two distinct ways: as countries where there have been few reported variants when discussing studies focused on specific genes, and as ethnicities with few studies or no representation to date in GWAS. *APP*,

*PSEN1* and *PSEN2* are the three genes harboring pathogenic mutations typically causing early-onset AD (EOAD) (age-at-onset below 65 years) in an autosomal dominant manner. Five to ten percent of EOAD is due to mutations in these genes. Over thirty loci have been significantly associated with risk for late-onset AD (LOAD) including, *ABCA7, BIN1, CLU, CR1, SORL1* and *APOE*. The E4 allele of *APOE* is currently the strongest genetic risk factor for LOAD ^1^. The heritability of AD is estimated to be 79% with this being higher for EOAD ^2^, however, the SNP heritability of LOAD (defined as the phenotypic variance attributable to common variants), is predicted to be approximately 31% ^3^. This suggests that a large proportion of AD heritability is still to be identified and could be due to rare variants.

Approximately 78% of all genetic studies from the NHGRI-EBI GWAS Catalog have been conducted on individuals of European origin ^4^. To note, the labels “European,” “non-Hispanic white” and “Caucasian” are commonly used interchangeably in the literature. In this review, we refer to these populations as described in the papers referenced and use the term non-Hispanic white (NHW) to broadly refer to both Americans and Europeans. Not only are there few studies on other populations, but many variants identified by large genetic studies on NHWs have not been replicated in other populations. This may be derived from lower sample sizes used in the replication experiments, or it may suggest that AD has a different genetic architecture, and perhaps even different mechanism(s) of disease in different populations. Although, it has been suggested that the same molecular pathways are disrupted in AD across populations, differences may occur in the genes involved and the specific impinging points in these pathways. Studying different populations is, thus, critical to our better understanding of the genetics of AD. This review aims to highlight our progress in studying AD across the world by discussing the key findings from such studies and corroborating insights and best practices to lead us throughout the next phase of human AD genetic studies.

### AD Mendelian genetics around the World

We first conducted a thorough literature search on 26th October 2019 by programmatically searching PubMed for the following terms: “Alzheimer* AND (PS1 OR PS2 OR PSEN1 OR APP OR PSEN2 OR presenilin OR amyloid precursor protein OR S182 OR E5-1) AND (mutation* or variant*) AND “+ country name. In order to interpret the 7,781 results, we filtered our results based on the metrics shown in Figure 1. From an initial 7,781 articles, we considered 377 studies. From this search, patients from 51 countries had variants reported in at least one of these 3 genes (Figure 2). Figure 3 shows the number of definitely and potentially pathogenic missense variants or indels in each gene reported in patients from each country from 377 articles. We used these results as the basis to determine how much AD genetics has been researched across populations. This approach assumes screening for Mendelian genes as an indication of research being conducted in a population. We have also attempted to consider other confounding factors as detailed below.

**Figure 1.**
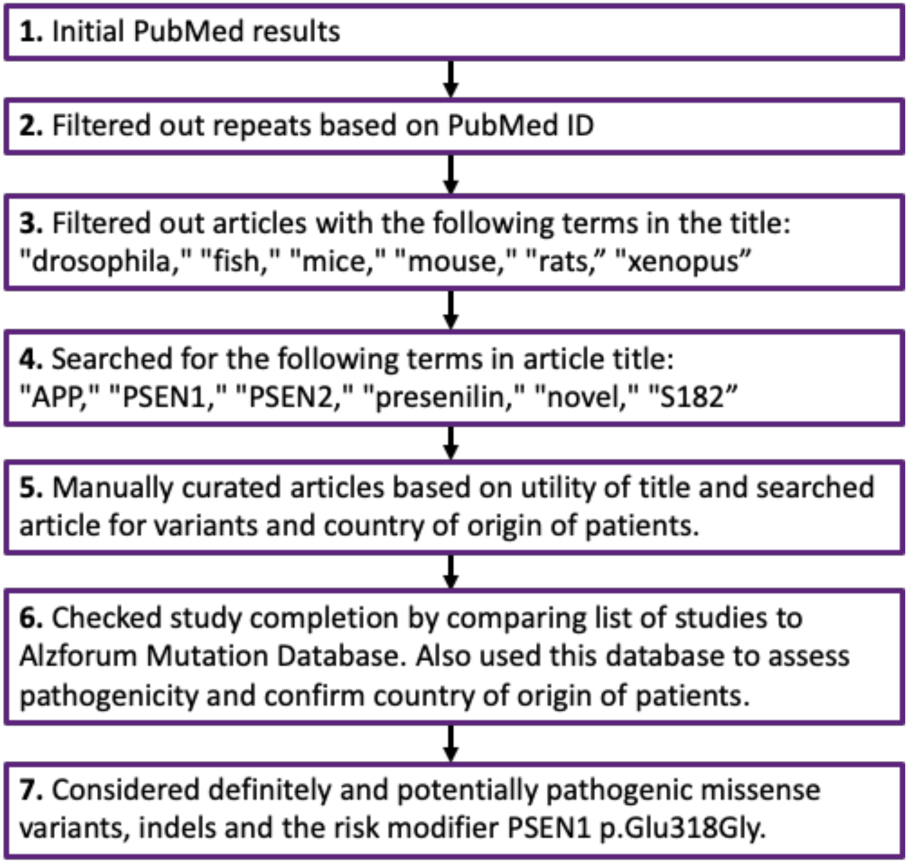
Flowchart to depict the steps taken for interpreting the literature on AD genetics collected through our PubMed search.

**Figure 2.**
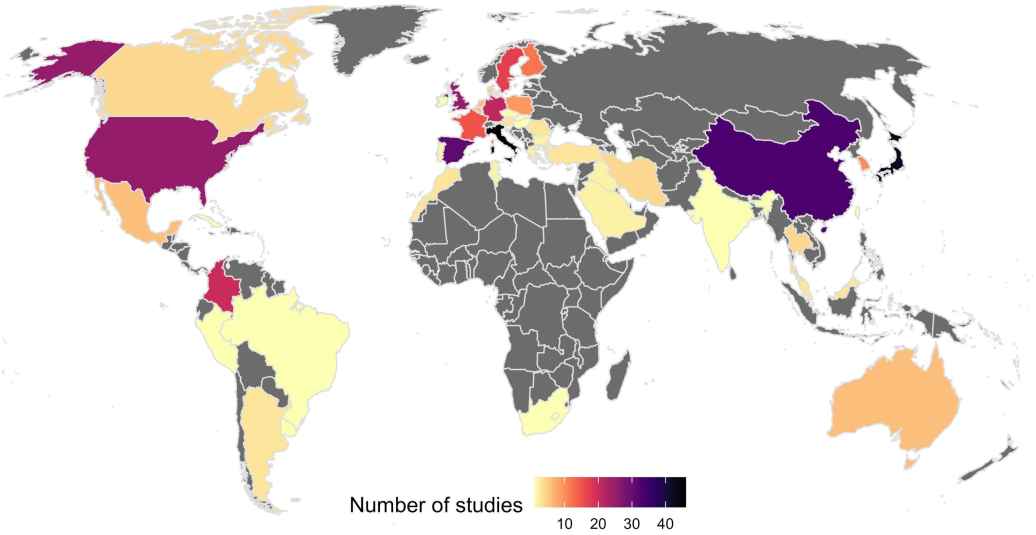
World map with number of studies reporting variants in *APP, PSEN1* or *PSEN2* in patients per country. Variants included are the risk modifier PSEN1:pGlu318Gly and missense and indels reported in AD cases as definitely or potentially pathogenic. Grey depicts countries with no studies on patients reporting variants in *APP, PSEN1* or *PSEN2.* The number of studies reported ranges from 1 study in countries depicted in yellow (Brazil, Cuba, Hungary, Israel, Peru, Serbia, Czech Republic, South Africa, Uruguay, Slovenia, Ireland, Slovakia, Taiwan and India) to 46 studies reported in Italy depicted in black, closely followed by 43 studies in Japan and 33 studies in China. References are in supplementary table 1.

**Figure 3.**
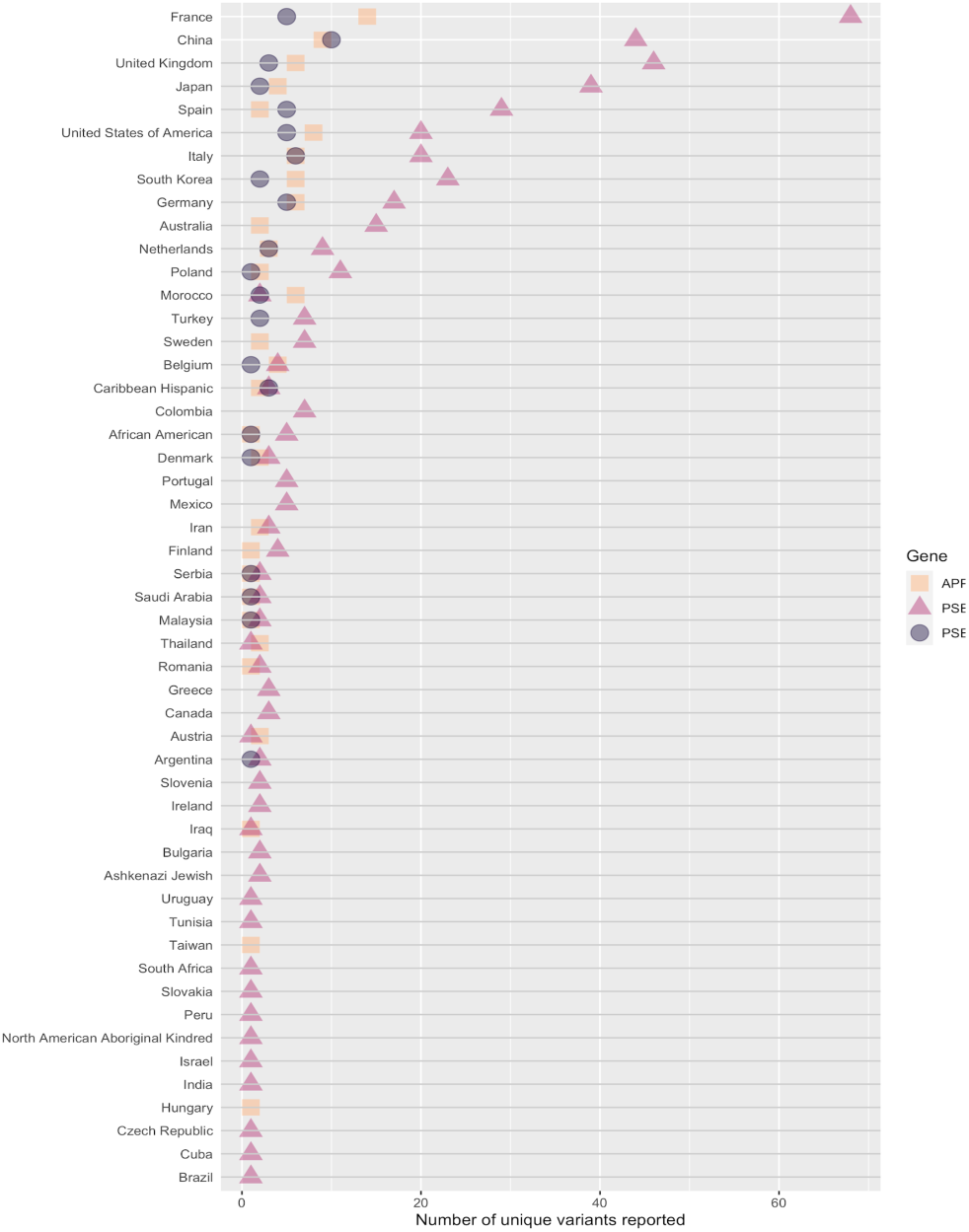
*APP, PSEN1* and *PSEN2* variants reported in patients from each country or population. Variants included are the risk modifier PSEN1:pGlu318Gly and missense and indels reported in AD cases as potentially pathogenic. We can clearly see that variants in *PSEN1* are the most common, with patients from the majority of countries reporting between 1-15 unique variants in this gene. No reports were found describing *PSEN1* variants in patients from Hungary or Taiwan, which is unusual given that pathogenic variants in this gene are the most frequent Mendelian cause of AD. This may be accounted for by the fact that, if patient country was not specifically reported, then study was not included. Patients from China, France, Italy, Japan, Korea, Spain, UK and US report over this range. Morocco stands out as having patients with more reports of *APP* variants compared to *PSEN1* variants; these are all frameshift variants and may not have been reported in other studies if they focused on missense, although, we can see from gnomAD that indels in *APP* are rare. In addition to patients from specific countries, there are also single-study reports of variants in patients from distinct populations (North American Aboriginal Kindred, Ashkenazi Jewish, African American and Caribbean Hispanic) which have also been included since these are distinct populations which encompass different countries. References are in supplementary table 1.

France is the country with the most variants reported, followed by China. When crossing the data derived from Figure 2 with that of Figure 3, it is clear that there are many more studies reporting few or individual variants in Chinese cohorts, as opposed to a small number of large studies in France reporting many variants. Many of the variants in the publications originating in China are novel and not replications of previous findings. Out of a total of 63 variants reported in China, 34 of these are unique to China since they have not been reported in any other countries to date (see supplementary table 2). It is interesting to note that the South of Europe has a considerable number of studies originating from Spain and Italy, but that Portugal is an understudied population for AD genetics. This is an important fact that can be missed if this population is combined with the Spanish and broadly referred to as Iberian.

In Colombia, two reported variants include: the PSEN1:p.Ile416Thr, which originated on an African haplotype ^5^, and the widely reported PSEN1:p.Glu280Ala, which segregates in a large community of over 5,000 people from Medellín, Colombia ^6^. In this understudied, yet highly informative population, it has been shown that elevated levels of tau deposition measured by PET imaging precede clinical onset of AD by approximately 6 years. This is important, since the ability to identify at-risk individuals prior to any symptom onset could be used to develop improved clinical trial designs for new therapies for AD. Accordingly, there is an ongoing phase II clinical trial (NCT01998841), involving presymptomatic carriers of the PSEN1:p.Glu280Ala, which tests the drug Crenezumab to target amyloid beta plaques. Even in such a unique population, composed of a very large number of individuals with the same mutation, and thus, very predictable disease course, positive results from this clinical trial could potentially impact patients across the whole world. In addition to the unique ability to develop genetically informed clinical trials in this extended family, the finding of the same mutation in such a large number of individuals also enables for the study of genetic modifiers of the disease. Recently it was found that the E2 allele of *APOE* delays the age at onset (AAO) of disease in cases carrying the PSEN1:p.Glu280Ala mutation ^7^, and the DAOA:p.Arg30Lys variant was found to be significantly associated (P=1.94×10^−4^) with AAO ^8^. More recently, one individual from this kindred demonstrated high resilience to AD with a delay of over 30 years on their AAO of MCI, despite high amyloid beta plaque load and carrying the PSEN1:p.Glu280Ala. They were identified as homozygous for the rare *APOE* E3 Christchurch variant p.Arg136Ser (equivalent to APOE:p.Arg154Ser) ^9^.

Additional reports suggest that the PSEN1 Glu280 residue is important in normal protein function for many populations. At the same protein residue, there is a report of another variant, PSEN1:p.Glu280Lys, in three siblings with EOAD from a Malaysian family ^10^. Ten additional family members (6 with neuropsychiatric symptoms and 4 asymptomatic) were screened and were negative for this variant, suggesting segregation with the disease in that family. Two additional *PSEN1* variants in apparent NHWs, PSEN1:p.Glu280Gly ^11^ and PSEN1:p.Glu280Gln ^12^, have also been reported in AD cases.

With about 65% protein sequence similarity to PSEN1, the highly homologous *PSEN2* gene also harbors mutations that cause EOAD. These two proteins are components of the gamma-secretase complex, which cleaves APP, and in addition to sequence, also share high structural similarity. Despite these similarities, mutations in *PSEN2* are rarer (59 in *PSEN1*, 36 in *PSEN2* per ClinVar assessed March 2020). The most common mutation in *PSEN2* is the p.Asn141Ile, first discovered in the Volga Germans. Interestingly, it was noted that seizures were reported in about 1/3 of Volga German cases with this *PSEN2* variant ^13,14^. This is intriguing and, in a similar way to the work described in the Colombian kindred above, opens the possibility of using the Volga population to identify genetic modifiers of AD that could be responsible for the seizures in this population. The cohort of 146 affected cases from 11 Volga German families, with seizures reported in 20 out of 64 of these cases, is a useful resource for such genotype-phenotype associations ^15^. Detecting if a patient is susceptible to developing seizures is important for close monitoring and prescription of medications.

Although there are no reports of variants in *APP, PSEN1* and *PSEN2* causing AD in Iceland, there is one report of an APP:p.Ala673Thr variant as protective against AD and cognitive decline in an elderly Icelandic population ^16^. This variant has since been identified in a Finnish individual who lived to 104.8 years ^17^. There are also reports of North Americans with this variant; some of whom are unaffected or have LOAD ^18^. All but one (who has Russian ancestry) of these reported individuals have broadly Scandinavian ancestry. To note, the frequency of this variant is much lower in populations outside Iceland and Finland, therefore, this protective variant would not have been identified without extremely large cohorts. These findings clearly exemplify the benefit of studying isolated and/or genetically distinct populations. The reduced genetic heterogeneity within such cohorts allows for the comparison of low-frequency variants between cases and controls; much larger sample sizes in populations with higher degrees of heterogeneity are required for sufficient detection power. It is interesting to note that at the same amino acid position in APP, there is one example of a recessive variant (APP:p.Ala673Val) which has been identified in two Italian siblings, one with early-onset dementia and the other with MCI; family members heterozygous for this variant were reported as unaffected ^19^.

The absence of pathogenic mutations in the three known AD genes in a given population can suggest different possibilities: 1) the absence of genetic testing in that population; 2) the possibility of other causative genes underlying the disease in that population; and/or 3) the presence of rare high risk genetic variants altering the susceptibility of individuals to AD in the population.

In conclusion, one can extrapolate if a population has been studied for AD genetics by searching for reports of variants in the genes known to harbor pathogenic variants that are implicated in Mendelian-inheritance of AD. The prediction of mutational damage to human genes should include population-specific analyses, as the predictions from current tools likely vary depending on ethnic background and the demographic history of the population ^20^. From our analysis, it is clear that there is wide variability not only in the number of studies across the globe, but also in the number of variants identified in each country. This has implications for future studies in the genetics of AD at a global level as it identifies populations to whom resources should be provided so that genetic studies can be performed. Even in the absence of effective therapies to prevent or delay AD, knowing the genetic status of an individual or family gives the opportunity for genetic counseling, informed preparation of life affairs (including reproductive options), and may contribute to achieve a more accurate in-life diagnosis ^21^.

### AD genetic risk in different populations

#### Representation of other populations in GWAS

We can use a similar approach to what was described in the previous section to determine which populations are lagging in terms of identifying common genetic variability associated with AD. To this end, and to explore the representation of different populations in AD GWAS, we used information from all available studies, with corresponding ancestry data, from the NHGRI-EBI GWAS catalog (16th January 2020 release). After filtering the studies for those focusing on AD, we identified 76 studies. In these, there was a clear overrepresentation of cohorts defined as broadly European (Figure 4). It is indisputable that large-scale genetic studies have revolutionized our understanding of the genetics of AD, however, this overrepresentation clearly highlights the need to study other populations in order for new genetic risk factors to be identified. To also emphasize this for diseases in general, the NHGRI-EBI GWAS catalog has released the GWAS Diversity Monitor which allows quasi-real-time monitoring of ancestries represented in GWAS studies ^22^.

**Figure 4.**
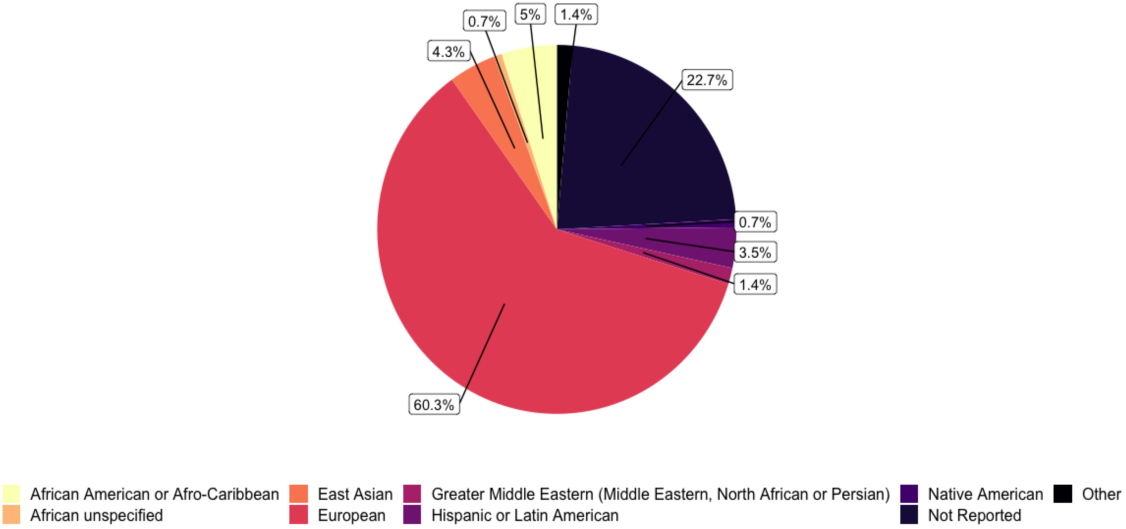
Broad ancestry represented in the 76 AD GWAS that are part of the NHGRI-EBI GWAS catalog. By separately including the different ancestries within each stage of a study, this plot provides a clearer view of the ancestries represented in AD GWAS to date. We can see that the representation of European ancestry is greater in AD GWAS compared to all GWAS ^4^. AD GWAS appear to have a greater representation of African ancestry compared to GWAS overall, but there is still a large percentage of studies with ancestry not reported. References and counts are in supplementary tables 3 and 4.

#### Translatability of known AD GWAS hits to date

In an attempt to compare the effect sizes of GWAS findings in NHW cohorts with other populations, we present ORs for the lead SNPs identified in the two most recent AD GWAS, which have also been tested in other populations (Table 1). We used the results presented by Kunkle et al. and Jansen et al. to represent NHW cohorts and searched the NHGRI-EBI GWAS catalog for studies of those same SNPs in other populations ^23,24^. We recorded 1) ORs from genome-wide significant SNPs and 2) ORs for variants in the same locus as these, from other populations (these variants did not have to reach genome-wide significance (GWS) to be recorded). In most cases, only the OR from the NHW data reached GWS, however there is one notable exception in *APOE*. The *APOE* E4 haplotype is the strongest genetic risk factor for LOAD and its dose-dependent effects were first discovered in 1993 ^25^. It was subsequently found that the *APOE* E2 isoform is protective against AD ^26^. The large increase in risk of LOAD conferred by the E4 allele in NHWs is not observed in the African American (AA) population where the risk conferred by this allele is smaller ^27^. This disparity has also been described in studies analyzing molecular biomarkers such as CSF total and phosphorylated tau in *APOE* E4-positive AAs AD cases ^27^. Many studies report *APOE* allele frequencies (AFs) in different populations and, compared to Europeans, both Africans and Oceanians have higher *APOE* E4 AFs, with Asians having the lowest AF ^28^. The decreased risk observed with *APOE* E4 in AAs is due to local genomic African ancestry, with an estimated OR=2.34, as opposed to AAs with local genomic European ancestry in which the OR is estimated to be 3.05 ^29^. A similar situation has also been shown in Puerto-Ricans (OR= 1.26 on African background, OR=4.49 on European background) ^29^ and may suggest that the Caribbean Hispanic GWAS AD cases have a stronger African background accounting for their lower OR when compared to other populations in Table 1. *APOE* E4 carrier frequency in South America/Mexico (57.3%) is reported to be similar to that of USA/Canada (55.8%) ^30^. This extends our understanding from *APOE* E4 or E2 in conferring risk or protection, respectively, to appreciating how population genetic background, and even ancestry local to the *APOE* locus, as defined by Rajabli et al., can modify such effects ^29^.

**Table 1.**
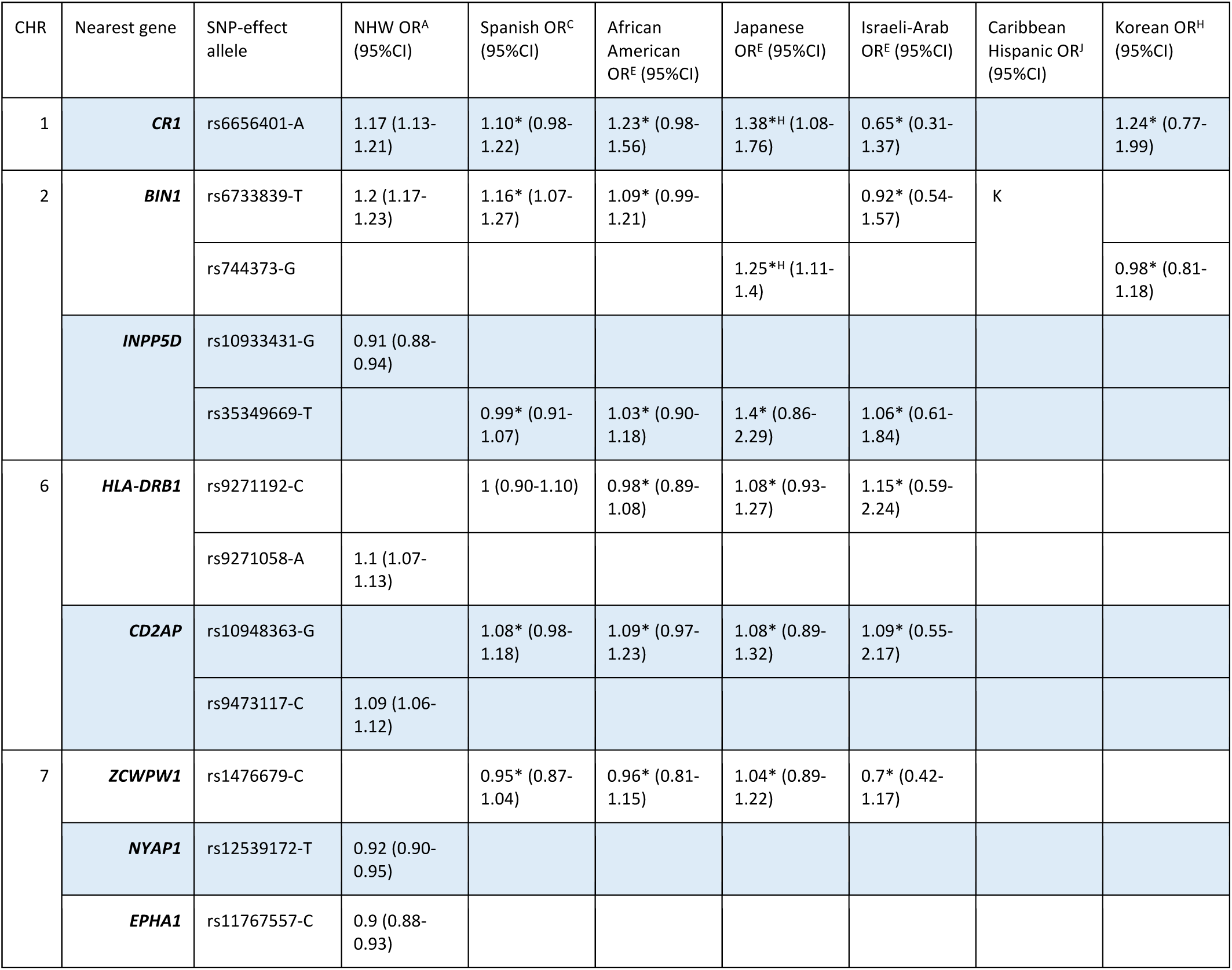
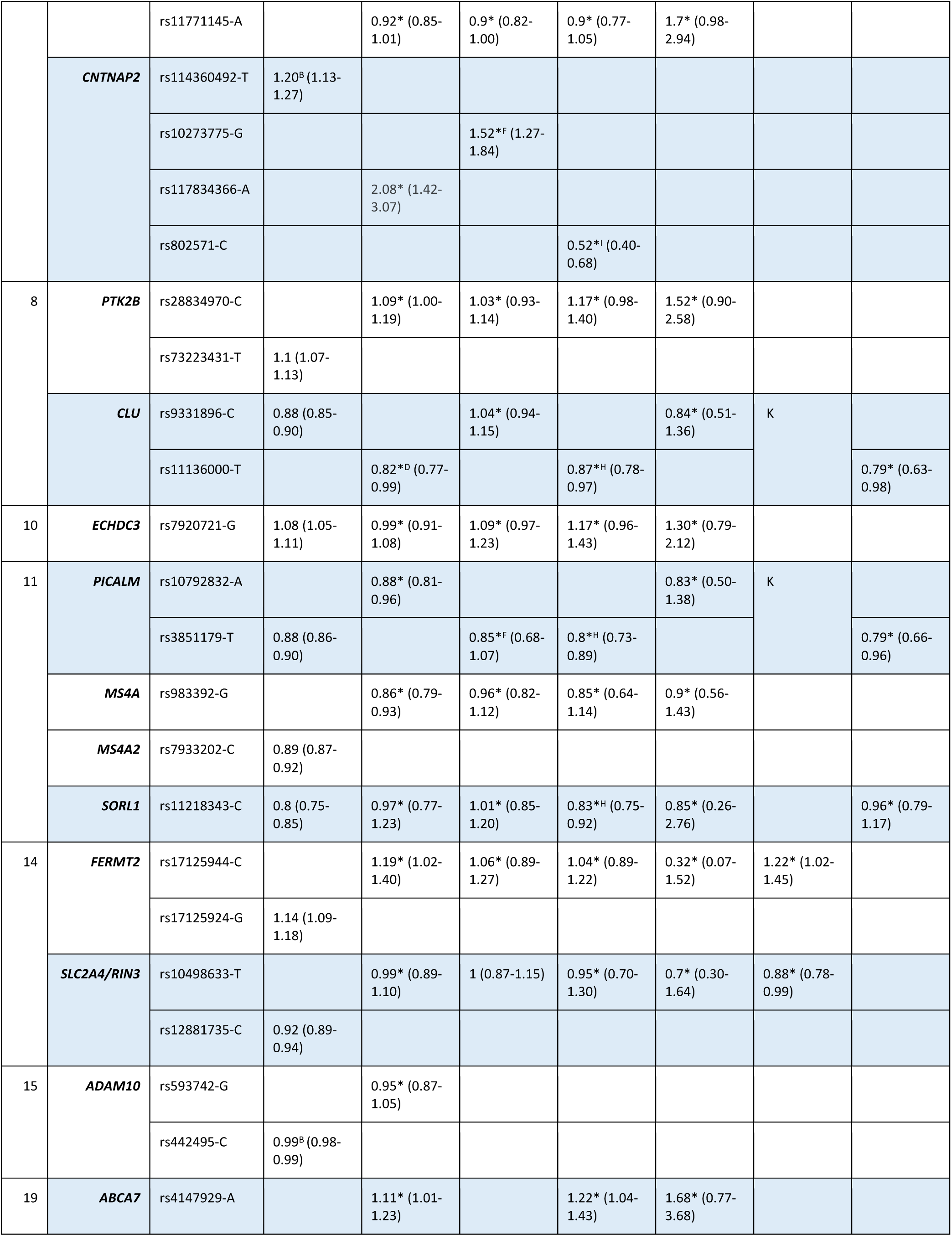
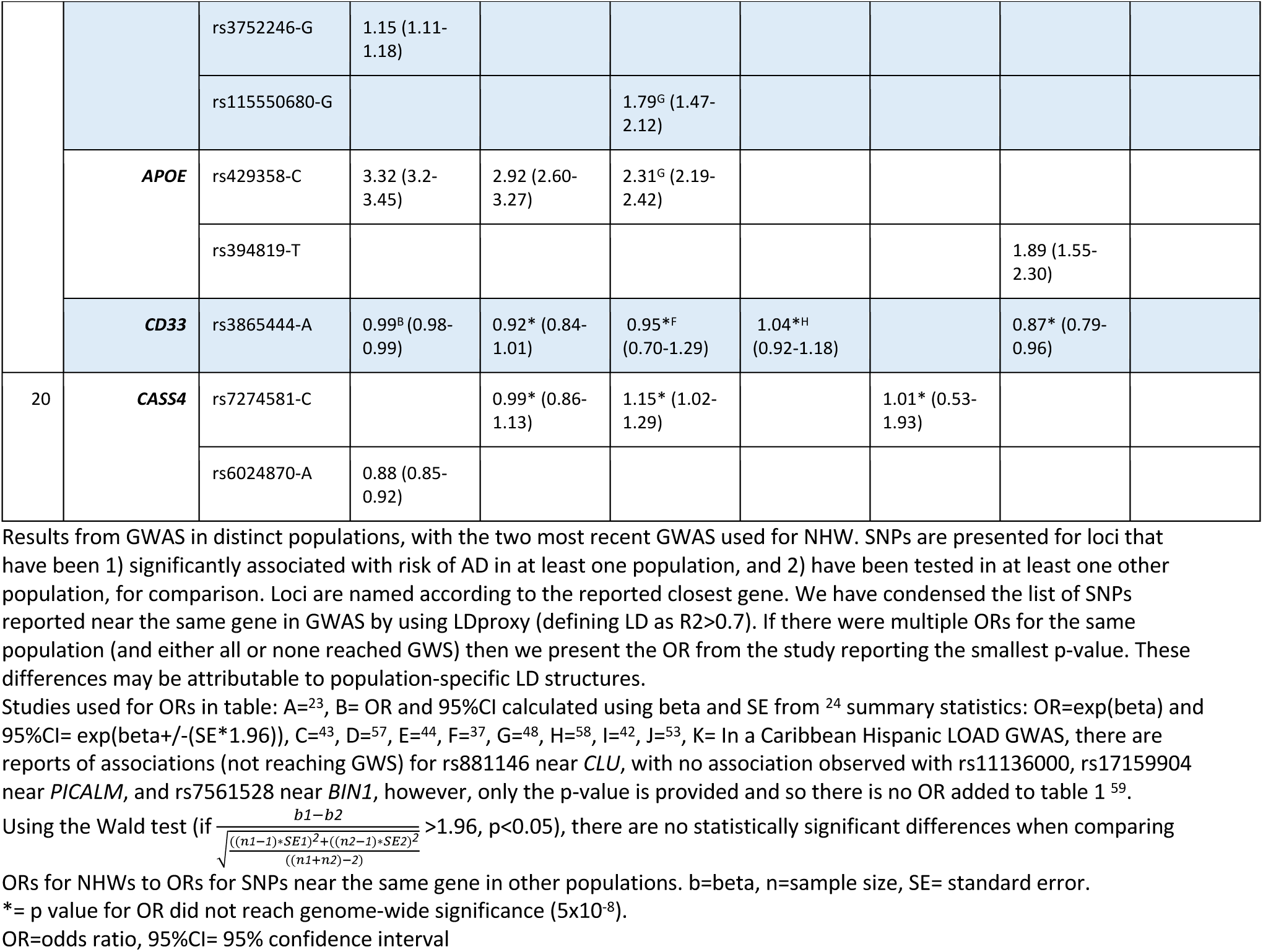
Odds ratios in different populations for GWAS hits that were significant in at least one population.

Although based on a small sample size, the data suggests that both *CR1* and *FERMT2* have an opposite direction of effect in Israeli-Arabs. It should be noted that, due to the sample size, there are very large confidence intervals associated with these ORs. Despite not always reaching GWS, the *CLU* rs9331896-C is associated with lower risk of AD in most populations, with the exception, with an OR close to 1, of African Americans. In line with this finding, Tycko et al., failed to identify any *CLU* polymorphisms associated with AD in African Americans ^31^. Moreover, a meta-analysis of 14 cohorts (total of 7,070 AD cases and 8,169 controls) concluded that rs11136000 in *CLU* is only associated with AD in NHWs (OR=0.91), and not in AA, Arab or Caribbean Hispanic populations ^32^. A study on a small South Indian cohort (243 cases and 164 controls) also reported no association between rs11136000 and AD, following genotyping of this polymorphism ^33^. In the Chinese population, conflicting reports describing meta-analyses have been published with some reporting a significant association between rs11136000 and LOAD ^34^, while others concluded that there was no effect ^35^. More recently, in a large meta-analysis including 24 individual studies, Almeida et al. showed rs11136000 to be protective; an association that remained statistically significant for the Caucasian as well as the mixed samples (from China, Japan, India, and Turkey). As expected, there were high levels of genetic heterogeneity in the mixed samples, highlighting the need to study associations in large enough cohorts of individual populations ^36^.

In the *BIN1* locus two SNPs have been reported with disparate associations in different populations, suggesting that independent signals exist in these populations (Table 1). The first, rs6733839, has been widely reported as a risk variant in various populations, however, in AAs there was no strong evidence of association ^37^. The second, rs744373, was identified as a risk variant in Japanese with no evidence for association in AA or Turkish populations ^38^. Following targeted sequencing and follow-up genotyping, Vardarajan et al., 2015 found BIN1:p.Lys358Arg (rs138047593) in 8 Caucasian and 6 Hispanic LOAD patients; segregation was shown in 2 out of 6 Caribbean Hispanic families ^39^. The authors reported higher frequency of this variant in familial cases (0.0859) where there was more than one affected individual in a family, compared to unaffected carriers (0.0084), and speculated that this increased *BIN1* rs138047593 frequency may be due to epistasis with other risk factors (genetic or environmental). Segregation was not tested in Caucasians because 7 out of 8 of them were from the Toronto LOAD study where there were no reported families ^39^. Although one Caucasian patient was from the NIA-LOAD study that includes families, segregation was not tested because the allele frequency of this variant for Caucasians in the ExAC database was similar to that observed in the LOAD cohort, suggesting that it was not associated with LOAD ^39^. Taken together these data show that *BIN1* variants have a similar effect in Asian and NHW populations ^40,41^, and that large Caribbean Hispanic families have enabled the finding of rare variants that seem to segregate with the disease.

Although there are no additional studies of the *EPHA1* lead GWAS SNPs in other populations, Vardarajan et al. reported segregation of the EPHA1:p.Pro460Leu variant (rs202178565) in 4 affected LOAD individuals and absence in three unaffected family members of a Caribbean Hispanic family from the Dominican Republic ^39^. The same variant showed no evidence of association in the Caucasian population studied ^39^. The authors suggested that rs202178565 may play a stronger role in AD risk in the Caribbean Hispanic population, however, further studies in larger datasets are needed to confirm this hypothesis.

At the *CNTNAP2* locus, Jansen et al found a significant signal for rs114360492 ^24^. This locus was previously reported in both AAs as well as in Japanese, however, the direction of effect was opposite in these two populations (risk and protection, respectively); it should be noted that these associations did not reach GWS in these populations ^37,42^. More recently, a GWAS in a Spanish cohort reported a significant association with rs117834366 near *CNTNAP2* in a vascular dementia sub-cohort at the discovery stage; this had a very high OR of 6.03 (3.22-11.2) ^43^. This finding indicated the possibility that the results observed by Jansen et al. could be due to the inclusion of vascular dementia cases in their AD cohort. However, the two variants show low frequency in the European population, which increases the likelihood of erroneous results due to low statistical power in smaller cohorts.

The *ECHDC3* locus was recently reported in NHWs ^23^. Interestingly, prior to this finding, the same locus had been reported in a transethnic GWAS where rs7920721-G reached significance when samples from all populations (European ancestry, AAs, Israeli-Arabs and Japanese) were combined ^44^. The significance was lost when populations were analyzed separately, although the direction of effect was maintained. These findings show the power of using transethnic approaches to identify novel loci when the effects are concordant.

Table 1 shows that *PICALM* rs10792832-A presented a protective effect in all populations tested with a GWAS approach. In both South and East Asians, rs3851179 is in LD (R^2^>0.7) with rs10792832 which we report in Table 1. However, when looking at studies that tested the specific variant outside of a GWAS framework, discordant results were found regarding association of rs3851179 and AD in the Asian population ^45,46^. It is plausible that this may be related to pooling Chinese and Japanese populations together to increase sample size, with small differences in genetic background leading to less clear results. It is also possible that *PICALM* variants may not affect both populations equally, since a recent study on the Han Chinese population reported no association between variants in *PICALM* and LOAD ^47^. Finally, in a South Indian population, there was no association found between *PICALM* rs3851179 and AD using a small cohort of 243 AD patients and 164 age-matched healthy controls ^33^.

The *ABCA7* locus is one of the strongest risk loci in AAs with rs115550680 showing an OR of 1.79 and reaching genome-wide significance ^48^. Although the authors mentioned that the SNP is in LD with the European risk variant (rs4147929), they have very different allele frequencies in both populations (leading to a high D’ and low R^2^). This, again, emphasizes the distinct genetic backgrounds among populations that may lead to population-specific genetic architectures of disease. These must be taken into consideration, for example, when translating the finding of risk alleles for PRS in different populations. Whether *ABCA7* plays a role in AD risk in the Asian is not clear with rs3764650 having been reported to be associated with AD in a Han Chinese population ^49^ and no association between *ABCA7* variants and AD being reported in two other studies on the same population ^50,51^.

A GWAS meta-analysis conducted using data from East Asian, North American and European populations (for a total of 31,106 cases and 55,653 controls) investigated the association of *CD33* rs3865444 with AD susceptibility ^52^. The authors reported no significant association in the East Asian population (Chinese, Japanese and Korean), however, an OR of 1.7 (p=0.0036) was reported for the major allele rs3865444-C in the Chinese subgroup with lower heterogeneity compared with the entire East Asian cohort ^52^. This is in line with the protective effect conferred by rs3865444-A in NHWs ^24^ and suggested in the Spanish, AAs and Caribbean Hispanics ^37,43,53^.

Many of the SNPs reaching significance in NHWs have been investigated in additional populations. When these are studied under a GWAS framework, the ORs in other populations usually do not reach significance, which is due, in all likelihood, to the smaller sample sizes being used. One example of this is the Israeli-Arab population where only 51 cases and 64 controls were used to calculate the ORs for the SNPs presented in Table 1. Given the clearly reduced statistical power in these smaller cohorts, results tend to focus on either a particular SNP or locus. This may account for the lack of ORs reported for Caribbean Hispanics in Table 1, compared to reports that we have mentioned. Nonetheless, key differences between populations can be observed for variants reaching significance, namely, the lower risk conferred by *APOE* in AAs, which is countered by the increased risk of *ABCA7*. A valuable tool in this space are transethnic GWAS, not only for comparison of ORs between populations, but also for aggregating enough samples to detect signals, which may then be replicated in other cohorts. The SNP near *ECHDC3* is a good example, where a novel significant association was initially reported in a transethnic GWAS before being replicated in the most recent GWAS in NHWs. Importantly, although this study aggregated samples of multiple ancestries, they also present results for separate populations, which enables future studies in these populations to build upon these population-specific results. Two studies could not be included in this review because they combined NHW samples with those of Caribbean Hispanic ancestry ^54^ and samples with Caucasian ancestry with AA samples ^55^. Lastly, it is important to note that post-GWAS analyses to investigate the potential effects of disease-associated SNPs are very informative and the field should gradually move its resources to systematically perform these for all nominated loci. The *CELF1/SPI1* locus is a great example of this, where rs10838725-C had been previously associated with increased risk of AD in NHWs (OR 1.08) and recent work has shown that *CELF1* rs1057233-G is associated with lower expression of *SPI1* in monocytes and macrophages, and delayed AD onset ^56^. It is important to test such associations in cells with different ancestral backgrounds; both the National Centralized Repository for Alzheimer’s Disease and Related Dementias and the Coriell Human Variation Panel may be a helpful resource with cell lines and DNA currently available from populations including African American, Middle Eastern, Italian and Greek.

#### Discovery of new AD risk variants in understudied populations

We performed a literature search to identify SNPs reaching GWS for association with AD in understudied populations (Table 2) and explored if these have been replicated in any subsequent studies. Since the original reports of these five associations, four of these have no reports of follow-up, highlighting a need to replicate these studies. A single study identified a 665kb duplication spanning *CACNA2D3* in post-mortem diagnosed LOAD from the Brain Bank of the Brazilian Aging Brain Study Group ^60^. Although this is not a confirmatory report, it is nonetheless interesting that a rare structural change in a post-mortem case overlapped a gene previously found to be associated with the same disease. There was also one report of statistically significant overexpression of *Cacna2d3* in the hypothalamus of aggressive male rats compared to domesticated rats ^61^. It would be interesting to study such changes in carriers of the risk SNP at this locus since some behavioral changes have also been reported in AD patients.

**Table 2.** Table of novel SNPs that were initially identified by reaching genome-wide significance in GWAS performed in understudied populations

| CHR | SNP-effect allele | Nearest gene | Population | Source |
| --- | --- | --- | --- | --- |
| 3 | rs7431992-A | *CACNA2D3* | Caribbean Hispanics using model A (logistic) | <sup>53</sup> |
| 5 | rs75002042-A | *FBXL7* | Caribbean Hispanics using model A (logistic) | <sup>53</sup> |
| 7 | rs112404845-T | *COBL* | AA- not GWS with logistic model but GWS with liability model with *APOE* E4 and *ABCA7* rs115550680 used as covariates | <sup>64</sup> |
| 13 | rs16961023-G | *SLC10A2* | AA- not GWS with logistic model but GWS with liability model with *APOE* E4 and *ABCA7* rs115550680 used as covariates | <sup>64</sup> |
| 19 | rs115553053-T | *HMHA1* | AA | <sup>48</sup> |

**Table 3.**
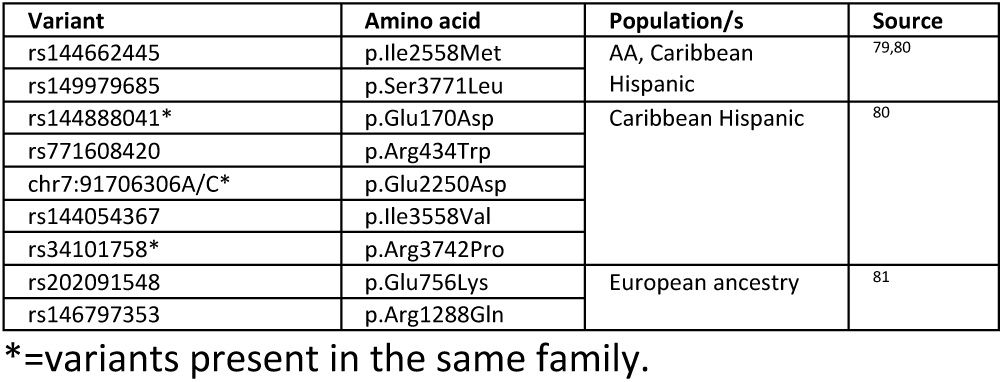
Variants in *AKAP9* reported in different populations.

Prior to the IGAP GWAS by Lambert et al., evidence for association at the *SORL1* locus had been obtained in a Japanese case-control study and subsequently replicated in a combined Japanese, Korean and Caucasian cohort ^58^. A protective effect seems to be present in other populations as well (Table 1), with the exception of AAs where, so far, there is no evidence of association. A study in a Caribbean Hispanic familial and sporadic LOAD cohort, found 17 exonic variants significantly associated with AD and 3 shown to segregate with disease in families under a dominant model ^62^.

An interesting understudied population is the Wadi Ara, where there is a high prevalence of AD, low frequency of *APOE* E4 and high degrees of consanguinity. In this population, Sherva et al. performed a GWAS and homozygosity mapping to identify new genes and risk loci for AD. Although they failed to find any significant SNPs in both the standard GWAS and the GWAS utilizing homozygous regions, they identified genes, such as *APBA1* or *AGER*, with AD-related biological functions, that were located within stretches of homozygosity nominally associated with AD ^63^. The lack of significant results likely stems from the fact that this was a study conducted in a small cohort of samples (124 cases and 142 controls). Focusing on the homozygous regions is an interesting approach to studying populations with high levels of consanguinity.

Since these associations were reported, there have been no further reports of associations between AD and *FBXL7, COBL, SLC10A2* or *HMHA1*.

GWAS mostly identify variants which convey small alterations in risk, whereas familial studies have the power to identify rare, high effect variants. The homozygous TREM2 variants p.Gln33X, p.Thr66Met and p.Tyr38Cys were first identified associated with dementia in a report of 3 separate Turkish FTD-like dementia families ^65^. These mutations, when homozygous, were previously known to cause polycystic lipomembranous osteodysplasia with sclerosing leukoencephalopathy (PLOSL), also known as Nasu Hakola Disease (NHD). The Turkish cases, however, displayed no apparent bone involvement in their phenotype with no history of joint pain or bone cysts. Since this initial finding in FTD-like families, Guerreiro et al., demonstrated that the heterozygous TREM2:p.Arg47His (rs75932628) is significantly associated with AD ^66^ - a result that was found independently in the Icelandic population ^67^ and has since been widely replicated, with significant associations reported for Spanish ^68,69^ and French populations ^70^. Conversely, there are also studies reporting no significant association between rs75932628 and AD in Japanese ^71^, African American ^72^, Chinese ^73–76^, and Iranian ^77^ populations, suggesting a population specific effect. In some populations not only is there no association, but also the p.Arg47His locus seems to be monomorphic, as is the case for the East Asian population in the latest version of gnomAD. Other variants in the gene have been identified in specific populations, such as the p.Gly55Arg variant reported in an Iranian cohort ^77^, and the p.Ala130Val in a Han Chinese LOAD patient ^73^. The noncoding variant rs7748513 is in LD with p.Arg47His and was reported to be associated with increased risk of AD in AAs ^78^. In addition to p.Arg47His, the p.Leu211Pro has also been shown to be associated with AD in AAs (p=0.01) ^72^.

Two rare variants in *AKAP9* were initially reported as nominally associated with AD in AAs ^79^. These two variants were also identified in individual Caribbean Hispanic LOAD cases ^80^. One of these two variants, the p.Arg434Trp, was reported in affected individuals from two large Caribbean Hispanic LOAD families ^80^. It is important to note that segregation was not shown for this variant, with absence in some affected individuals and presence in potentially presymptomatic individuals who were younger than the usual AAO of other family members. There was one exception of a single family member aged 78 who was unaffected but identified as a heterozygous carrier ^80^. Taken together this data argues against a fully penetrant mode for *AKAP9* mutations in this Caribbean Hispanic family. Moreover, 4 other *AKAP9* variants were reported in Caribbean Hispanic families, with 3 of them present in the same family ^80^. Two different *AKAP9* variants have subsequently been reported in LOAD families of European ancestry, although segregation was not shown for either of them ^81^.

Association of *CASP7* with AD was first reported in a Caribbean Hispanic haplotype association study ^82^. Since this report, a loss-of-function variant (rs10553596) has been reported to reduce the risk of AD in *APOE* E4 homozygotes ^83^. This haplotype association study also reported significant association between *LRP1B, TNFRSF1A, CDH1*, and *TG*, with AD susceptibility in Caribbean Hispanic individuals ^82^. Finally, the *CASP7* rs116437863 variant has been significantly associated with familial LOAD in individuals of European ancestry ^84^.

*NOTCH3* variants have been associated with AD risk relatively recently and thus the impact of such variants is still not entirely clear. The NOTCH3:p.Arg1231Cys (rs201680145-A) mutation was first reported in one Caucasian patient with cerebral autosomal dominant arteriopathy with subcortical infarcts and leukoencephalopathy (CADASIL) ^85^. There have since been several reports of this mutation in CADASIL patients from central Italy ^86^, in one Russian CADASIL patient (a compound heterozygote with NOTCH3:p.Phe984Cys) ^87^ and in a Turkish AD case ^88^. This variant is not reported in unrelated Chinese cases diagnosed with CADASIL ^89^. Since the report linking *NOTCH3* mutations with AD, there has been growing evidence supporting this association. Following exome sequencing, a missense *NOTCH3* variant (rs149307620, p.Ala284Thr) was reported in 10 AD cases (confirmed in 8), and not present in controls of European ancestry ^90^. Screening of WGS from the ADSP and ADNI datasets revealed this variant in one AD case, one MCI case and no controls. Moreover, two half-first cousins from a Utah high-risk pedigree both had rare missense variants in *NOTCH3* which were not identified in the WGS samples from the ADSP and ADNI datasets. A study on the Han Chinese population reported the NOTCH3:p.Glu585Ala in 1 probable AD case (out of 210 AD cases) with early onset familial AD, which was absent from 160 population matched controls ^91^. Despite these results, studies in more populations and in larger sample sizes are required to fully establish the role of *NOTCH3* variants in AD.

In summary, several variants and genes were initially identified and associated with AD in understudied populations that were later found to also have an effect in NHW individuals. This highlights the importance of performing family and cohort studies in diverse populations.

## Discussion

The human reference genome is not representative of an average genome for all ethnicities, instead it is largely representative of NHW ancestry only. As our understanding of the genetics of human disease continues to evolve, this creates important limitations. It is also possible that regions encompassing damaging variants in some populations are not mapped in the current reference genome and thus will not be detected. Efforts have been made to create ethnicity-specific reference genomes and there is some interest in designing a pan-genome to account for all genetic variability. Recent work to create a pan-genome of African individuals highlighted that there is over 296.5Mb of DNA that is not accounted for in the current human genome reference ^92^.

There are disparities in gene variants associated with AD between populations, both from an effect size as well as direction of effect perspectives. These may be due to early human migration out of Africa and subsequent founder effects. In AD, as in all other complex diseases, substantially more samples of NHW ancestry are used for initial discovery in genetic studies for all diseases. Interestingly, with regards to populations used for replication studies, samples of Asian ancestry are currently the most commonly used after NHW samples ^93^. The smaller sample sizes of AD cases obtained for other populations compared to NHWs, may be related to stigmatization of AD in some cultures, but also to funding availability for large-scale genetic research in those populations. However, it was recently shown that between 1990 and 2016 there has been an increase in both prevalence and burden of dementias in Eastern Mediterranean region countries ^94^, which may be due to a range of factors, such as an aging population or improved diagnostic methodologies. Based both on key past discoveries, and the potential for future findings, funding to facilitate the study of understudied populations is imperative. The immediate need to address ethnic and racial disparities in AD and related dementias has been the focus of a recent white paper ^95^, which developed a series of recommendations for future strategies. These included targeted study recruitment and retention of diverse ethnoracial populations, generalization of instruments and analytic methods across research groups, ensuring that ethnoracial sub-group data is reported, creation of methods to reduce impact of small sample size on observations, and development of statistical models of risk and protective factors relevant to ethnoracial groups, to better refine prevalence and incidence ^95^.

Data from the UK Biobank, which is largely European-based, has been used in AD GWAS by taking advantage of proximal AD status, where an individual is considered as AD-by-proxy if they report AD in one of their parents, while individuals with parents without history of AD are considered by-proxy-controls ^24,96^. This is an approach that could be used to increase the sample sizes of understudied populations and thus the power for detecting associations. As detailed above, it is critically important to have a better understanding of the allele frequencies across populations, something that population variant databases such as ExAC and gnomAD have enabled us to do, to a certain degree, for some populations. For Middle Eastern populations, at a smaller scale and with no information on family history, one available resource is the Greater Middle East Variome (WES data on 1,111 unrelated subjects from populations including Northwest and Northeast Africa, the Turkish Peninsula, the Syrian Desert, the Arabian Peninsula, and Persia and Pakistan) ^97^. Other population-specific databases include the GenomeAsia 100K Project, the Han Chinese genome database, the Japanese Human Genetic Variation Database and the Genome of the Netherlands ^98–101^. Additionally, studies on populations with large genetic bottlenecks, such as Finland, where there is increased frequency of extremely rare alleles, can increase the power for finding associations, without needing to drastically increase sample size ^102^.

Large cohorts with the same mutation, such as the Colombian, have demonstrated the power to study genetic disease modifiers, clinical biomarkers and therapeutics with the potential to translate findings to other populations. Related to the clinical setting, polygenic risk scores take into account SNPs across the genome that were associated with modulating risk for disease. The best predictors of risk are derived from the largest GWAS as these will be the best powered studies to identify risk loci and estimate their effect size. Currently, the largest GWAS are on NHWs, which raises the question of how valid these PRSs are for prediction of AD risk in other populations. If greater diversity is not prioritized in genetic studies, current PRSs may be irrelevant for the majority of the world’s populations and may exacerbate health disparities ^103^.

The vast majority of genetic findings in complex disease in general and AD in particular have been made in broadly NHW populations. We have highlighted here how important findings from other populations have been reported over the years and how they have improved our understanding of the genetic bases of this disease. A shift in funding strategies and a broader collaborative mindset is required to allow us to create the next step-change that is needed to move the field to a complete understanding of the genetic architecture of AD across populations.

## Author Contributions and Notes

We contributed equally to the preparation and writing of the manuscript, and we all approved the final version.

## Acknowledgements

We thank the Van Andel Institute Bioinformatics and Biostatistics Core, especially Emily Wolfrum and Zach Madaj, for their statistical expertise. Research reported in this publication was supported by the National Institute on Aging of the National Institutes of Health under Award Number R01AG067426. The content is solely the responsibility of the authors and does not necessarily represent the official views of the National Institutes of Health.

## References

1. Guerreiro R, Hardy J. Genetics of Alzheimer’s disease. Neurotherapeutics. 2014;11(4):732–737.

2. Gatz M, Reynolds CA, Fratiglioni L, et al. Role of genes and environments for explaining Alzheimer disease. Arch Gen Psychiatry. 2006;63(2):168–174.

3. Ridge PG, Hoyt KB, Boehme K, et al. Assessment of the genetic variance of late-onset Alzheimer’s disease. Neurobiol Aging. 2016;41:200.e13-e200.e20.

4. Morales J, Welter D, Bowler EH, et al. A standardized framework for representation of ancestry data in genomics studies, with application to the NHGRI-EBI GWAS Catalog. Genome Biol. 2018;19(1):21.

5. Ramirez Aguilar L, Acosta-Uribe J, Giraldo MM, et al. Genetic origin of a large family with a novel PSEN1 mutation (Ile416Thr). Alzheimers Dement. 2019;15(5):709–719.

6. Lopera F, Ardilla A, Martínez A, et al. Clinical features of early-onset Alzheimer disease in a large kindred with an E280A presenilin-1 mutation. JAMA. 1997;277(10):793–799.

7. Vélez JI, Lopera F, Sepulveda-Falla D, et al. APOE*E2 allele delays age of onset in PSEN1 E280A Alzheimer’s disease. Mol Psychiatry. 2016;21(7):916–924.

8. Vélez JI, Rivera D, Mastronardi CA, et al. A Mutation in DAOA Modifies the Age of Onset in PSEN1 E280A Alzheimer’s Disease. Neural Plast. 2016;2016:9760314.

9. Arboleda-Velasquez JF, Lopera F, O’Hare M, et al. Resistance to autosomal dominant Alzheimer’s disease in an APOE3 Christchurch homozygote: a case report. Nat Med. 2019;25(11):1680–1683.

10. Ch’ng G-S, An SSA, Bae SO, Bagyinszky E, Kim S. Identification of two novel mutations, PSEN1 E280K and PRNP G127S, in a Malaysian family. Neuropsychiatr Dis Treat. 2015;11:2315–2322.

11. Alzheimer’s Disease Collaborative Group. The structure of the presenilin 1 (S182) gene and identification of six novel mutations in early onset AD families. Nat Genet. 1995;11(2):219–222.

12. Rogaeva E, Bergeron C, Sato C, et al. PS1 Alzheimer’s disease family with spastic paraplegia: the search for a gene modifier. Neurology. 2003;61(7):1005–1007.

13. Levy-Lahad E, Wasco W, Poorkaj P, et al. Candidate gene for the chromosome 1 familial Alzheimer’s disease locus. Science. 1995;269(5226):973–977.

14. Rogaev EI, Sherrington R, Rogaeva EA, et al. Familial Alzheimer’s disease in kindreds with missense mutations in a gene on chromosome 1 related to the Alzheimer’s disease type 3 gene. Nature. 1995;376(6543):775–778.

15. Jayadev S, Leverenz JB, Steinbart E, et al. Alzheimer’s disease phenotypes and genotypes associated with mutations in presenilin 2. Brain. 2010;133(Pt 4):1143–1154.

16. Jonsson T, Atwal JK, Steinberg S, et al. A mutation in APP protects against Alzheimer’s disease and age-related cognitive decline. Nature. 2012;488(7409):96–99.

17. Kero M, Paetau A, Polvikoski T, et al. Amyloid precursor protein (APP) A673T mutation in the elderly Finnish population. Neurobiol Aging. 2013;34(5):1518.e1-e3.

18. Wang L-S, Naj AC, Graham RR, et al. Rarity of the Alzheimer disease-protective APP A673T variant in the United States. JAMA Neurol. 2015;72(2):209–216.

19. Di Fede G, Catania M, Morbin M, et al. A recessive mutation in the APP gene with dominant-negative effect on amyloidogenesis. Science. 2009;323(5920):1473–1477.

20. Hussin JG, Hodgkinson A, Idaghdour Y, et al. Recombination affects accumulation of damaging and disease-associated mutations in human populations. Nat Genet. 2015;47(4):400–404.

21. Cohn-Hokke PE, Elting MW, Pijnenburg YAL, van Swieten JC. Genetics of dementia: update and guidelines for the clinician. Am J Med Genet B Neuropsychiatr Genet. 2012;159(6):628–643.

22. Mills MC, Rahal C. The GWAS Diversity Monitor tracks diversity by disease in real time. Nat Genet. 2020;52(3):242–243.

23. Kunkle BW, Grenier-Boley B, Sims R, et al. Genetic meta-analysis of diagnosed Alzheimer’s disease identifies new risk loci and implicates Aβ, tau, immunity and lipid processing. Nat Genet. 2019;51(3):414–430.

24. Jansen IE, Savage JE, Watanabe K, et al. Genome-wide meta-analysis identifies new loci and functional pathways influencing Alzheimer’s disease risk. Nat Genet. 2019;51(3):404–413.

25. Corder EH, Saunders AM, Strittmatter WJ, et al. Gene dose of apolipoprotein E type 4 allele and the risk of Alzheimer’s disease in late onset families. Science. 1993;261(5123):921–923.

26. Corder EH, Saunders AM, Risch NJ, et al. Protective effect of apolipoprotein E type 2 allele for late onset Alzheimer disease. Nat Genet. 1994;7(2):180–184.

27. Morris JC, Schindler SE, McCue LM, et al. Assessment of Racial Disparities in Biomarkers for Alzheimer Disease. JAMA Neurol. 2019;76(3):264–273.

28. Kamboh MI. Apolipoprotein E polymorphism and susceptibility to Alzheimer’s disease. Hum Biol. 1995;67(2):195–215.

29. Rajabli F, Feliciano BE, Celis K, et al. Ancestral origin of ApoE ε4 Alzheimer disease risk in Puerto Rican and African American populations. PLoS Genet. 2018;14(12):e1007791.

30. Cummings JL, Atri A, Ballard C, et al. Insights into globalization: comparison of patient characteristics and disease progression among geographic regions in a multinational Alzheimer’s disease clinical program. Alzheimers Res Ther. 2018;10(1):116.

31. Tycko B, Feng L, Nguyen L, et al. Polymorphisms in the human apolipoprotein-J/clusterin gene: ethnic variation and distribution in Alzheimer’s disease. Hum Genet. 1996;98(4):430–436.

32. Jun G, Naj AC, Beecham GW, et al. Meta-analysis confirms CR1, CLU, and PICALM as alzheimer disease risk loci and reveals interactions with APOE genotypes. Arch Neurol. 2010;67(12):1473–1484.

33. Shankarappa BM, Kota LN, Purushottam M, et al. Effect of CLU and PICALM polymorphisms on AD risk: A study from south India. Asian J Psychiatr. 2017;27:7–11.

34. Ma J-F, Liu L-H, Zhang Y, et al. Association study of clusterin polymorphism rs11136000 with late onset Alzheimer’s disease in Chinese Han population. Am J Alzheimers Dis Other Demen. 2011;26(8):627–630.

35. Han Z, Qu J, Zhao J, Zou X. Analyzing 74,248 Samples Confirms the Association Between CLU rs11136000 Polymorphism and Alzheimer’s Disease in Caucasian But Not Chinese population. Sci Rep. 2018;8(1):11062.

36. Almeida JFF, Dos Santos LR, Trancozo M, de Paula F. Updated Meta-Analysis of BIN1, CR1, MS4A6A, CLU, and ABCA7 Variants in Alzheimer’s Disease. J Mol Neurosci. 2018;64(3):471–477.

37. Logue MW, Schu M, Vardarajan BN, et al. A comprehensive genetic association study of Alzheimer disease in African Americans. Arch Neurol. 2011;68(12):1569–1579.

38. Kaya G, Gündüz E, Acar M, et al. Potential genetic biomarkers in the early diagnosis of Alzheimer disease: APOE and BIN1. Turk J Med Sci. 2015;45(5):1058–1072.

39. Vardarajan BN, Ghani M, Kahn A, et al. Rare coding mutations identified by sequencing of Alzheimer disease genome-wide association studies loci. Ann Neurol. 2015;78(3):487–498.

40. Liu G, Zhang S, Cai Z, et al. BIN1 gene rs744373 polymorphism contributes to Alzheimer’s disease in East Asian population. Neurosci Lett. 2013;544:47–51.

41. Zhu R, Liu X, He Z. The Bridging Integrator 1 Gene Polymorphism rs744373 and the Risk of Alzheimer’s Disease in Caucasian and Asian Populations: An Updated Meta-Analysis. Mol Neurobiol. 2017;54(2):1419–1428.

42. Hirano A, Ohara T, Takahashi A, et al. A genome-wide association study of late-onset Alzheimer’s disease in a Japanese population. Psychiatr Genet. 2015;25(4):139–146.

43. Moreno-Grau S, de Rojas I, Hernández I, et al. Genome-wide association analysis of dementia and its clinical endophenotypes reveal novel loci associated with Alzheimer’s disease and three causality networks: The GR@ACE project. Alzheimers Dement. 2019;15(10):1333–1347.

44. Jun GR, Chung J, Mez J, et al. Transethnic genome-wide scan identifies novel Alzheimer’s disease loci. Alzheimers Dement. 2017;13(7):727–738.

45. Liu G, Zhang S, Cai Z, et al. PICALM gene rs3851179 polymorphism contributes to Alzheimer’s disease in an Asian population. Neuromolecular Med. 2013;15(2):384–388.

46. Liu G, Zhang L, Feng R, et al. Lack of association between PICALM rs3851179 polymorphism and Alzheimer’s disease in Chinese population and APOEε4-negative subgroup. Neurobiol Aging. 2013;34(4):1310.e9-e10.

47. Jiang T, Yu J-T, Tan M-S, et al. Genetic variation in PICALM and Alzheimer’s disease risk in Han Chinese. Neurobiol Aging. 2014;35(4):934.e1-e3.

48. Reitz C, Jun G, Naj A, et al. Variants in the ATP-binding cassette transporter (ABCA7), apolipoprotein E ϵ4,and the risk of late-onset Alzheimer disease in African Americans. JAMA. 2013;309(14):1483–1492.

49. Liao Y-C, Lee W-J, Hwang J-P, et al. ABCA7 gene and the risk of Alzheimer’s disease in Han Chinese in Taiwan. Neurobiol Aging. 2014;35(10):2423.e7-e2423.e13.

50. Liu L-H, Xu J, Deng Y-L, et al. A complex association of ABCA7 genotypes with sporadic Alzheimer disease in Chinese Han population. Alzheimer Dis Assoc Disord. 2014;28(2):141–144.

51. Tan L, Yu J-T, Zhang W, et al. Association of GWAS-linked loci with late-onset Alzheimer’s disease in a northern Han Chinese population. Alzheimers Dement. 2013;9(5):546–553.

52. Li X, Shen N, Zhang S, et al. CD33 rs3865444 Polymorphism Contributes to Alzheimer’s Disease Susceptibility in Chinese, European, and North American Populations. Mol Neurobiol. 2015;52(1):414–421.

53. Tosto G, Fu H, Vardarajan BN, et al. F-box/LRR-repeat protein 7 is genetically associated with Alzheimer’s disease. Ann Clin Transl Neurol. 2015;2(8):810–820.

54. Wijsman EM, Pankratz ND, Choi Y, et al. Genome-wide association of familial late-onset Alzheimer’s disease replicates BIN1 and CLU and nominates CUGBP2 in interaction with APOE. PLoS Genet. 2011;7(2):e1001308.

55. Hollingworth P, Sweet R, Sims R, et al. Genome-wide association study of Alzheimer’s disease with psychotic symptoms. Mol Psychiatry. 2012;17(12):1316–1327.

56. Huang K-L, Marcora E, Pimenova AA, et al. A common haplotype lowers PU.1 expression in myeloid cells and delays onset of Alzheimer’s disease. Nat Neurosci. 2017;20(8):1052–1061.

57. Seshadri S, Fitzpatrick AL, Ikram MA, et al. Genome-wide analysis of genetic loci associated with Alzheimer disease. JAMA. 2010;303(18):1832–1840.

58. Miyashita A, Koike A, Jun G, et al. SORL1 is genetically associated with late-onset Alzheimer’s disease in Japanese, Koreans and Caucasians. PLoS One. 2013;8(4):e58618.

59. Lee JH, Cheng R, Barral S, et al. Identification of novel loci for Alzheimer disease and replication of CLU, PICALM, and BIN1 in Caribbean Hispanic individuals. Arch Neurol. 2011;68(3):320–328.

60. Villela D, Suemoto CK, Pasqualucci CA, Grinberg LT, Rosenberg C. Do Copy Number Changes in CACNA2D2, CACNA2D3, and CACNA1D Constitute a Predisposing Risk Factor for Alzheimer’s Disease? Front Genet. 2016;7:107.

61. Oshchepkov D, Ponomarenko M, Klimova N, et al. A Rat Model of Human Behavior Provides Evidence of Natural Selection Against Underexpression of Aggressiveness-Related Genes in Humans. Front Genet. 2019;10:1267.

62. Vardarajan BN, Zhang Y, Lee JH, et al. Coding mutations in SORL1 and Alzheimer disease. Ann Neurol. 2015;77(2):215–227.

63. Sherva R, Baldwin CT, Inzelberg R, et al. Identification of novel candidate genes for Alzheimer’s disease by autozygosity mapping using genome wide SNP data. J Alzheimers Dis. 2011;23(2):349–359.

64. Mez J, Chung J, Jun G, et al. Two novel loci, COBL and SLC10A2, for Alzheimer’s disease in African Americans. Alzheimers Dement. 2017;13(2):119–129.

65. Guerreiro RJ, Lohmann E, Brás JM, et al. Using exome sequencing to reveal mutations in TREM2 presenting as a frontotemporal dementia-like syndrome without bone involvement. JAMA Neurol. 2013;70(1):78–84.

66. Guerreiro R, Wojtas A, Bras J, et al. TREM2 variants in Alzheimer’s disease. N Engl J Med. 2013;368(2):117–127.

67. Jonsson T, Stefansson H, Steinberg S, et al. Variant of TREM2 associated with the risk of Alzheimer’s disease. N Engl J Med. 2013;368(2):107–116.

68. Benitez BA, Cooper B, Pastor P, et al. TREM2 is associated with the risk of Alzheimer’s disease in Spanish population. Neurobiol Aging. 2013;34(6):1711.e15-e17.

69. Ruiz A, Dols-Icardo O, Bullido MJ, et al. Assessing the role of the TREM2 p.R47H variant as a risk factor for Alzheimer’s disease and frontotemporal dementia. Neurobiol Aging. 2014;35(2):444.e1-e4.

70. Pottier C, Wallon D, Rousseau S, et al. TREM2 R47H variant as a risk factor for early-onset Alzheimer’s disease. J Alzheimers Dis. 2013;35(1):45–49.

71. Miyashita A, Wen Y, Kitamura N, et al. Lack of genetic association between TREM2 and late-onset Alzheimer’s disease in a Japanese population. J Alzheimers Dis. 2014;41(4):1031–1038.

72. Jin SC, Carrasquillo MM, Benitez BA, et al. TREM2 is associated with increased risk for Alzheimer’s disease in African Americans. Mol Neurodegener. 2015;10:19.

73. Jiao B, Liu X, Tang B, et al. Investigation of TREM2, PLD3, and UNC5C variants in patients with Alzheimer’s disease from mainland China. Neurobiol Aging. 2014;35(10):2422.e9-e2422.e11.

74. Ma J, Zhou Y, Xu J, et al. Association study of TREM2 polymorphism rs75932628 with late-onset Alzheimer’s disease in Chinese Han population. Neurol Res. 2014;36(10):894–896.

75. Wang P, Guo Q, Zhou Y, et al. Lack of association between triggering receptor expressed on myeloid cells 2 polymorphism rs75932628 and late-onset Alzheimer’s disease in a Chinese Han population. Psychiatr Genet. 2018;28(1):16–18.

76. Yu J-T, Jiang T, Wang Y-L, et al. Triggering receptor expressed on myeloid cells 2 variant is rare in late-onset Alzheimer’s disease in Han Chinese individuals. Neurobiol Aging. 2014;35(4):937.e1-e3.

77. Mehrjoo Z, Najmabadi A, Abedini SS, et al. Association Study of the TREM2 Gene and Identification of a Novel Variant in Exon 2 in Iranian Patients with Late-Onset Alzheimer’s Disease. Med Princ Pract. 2015;24(4):351–354.

78. Reitz C, Mayeux R, Alzheimer’s Disease Genetics Consortium. TREM2 and neurodegenerative disease. N Engl J Med. 2013;369(16):1564–1565.

79. Logue MW, Schu M, Vardarajan BN, et al. Two rare AKAP9 variants are associated with Alzheimer’s disease in African Americans. Alzheimers Dement. 2014;10(6):609-618.e11.

80. Vardarajan BN, Barral S, Jaworski J, et al. Whole genome sequencing of Caribbean Hispanic families with late-onset Alzheimer’s disease. Ann Clin Transl Neurol. 2018;5(4):406–417.

81. Cukier HN, Kunkle BK, Hamilton KL, et al. Exome Sequencing of Extended Families with Alzheimer’s Disease Identifies Novel Genes Implicated in Cell Immunity and Neuronal Function. J Alzheimers Dis Parkinsonism. 2017;7(4). doi: 10.4172/2161-0460.1000355

82. Shang Z, Lv H, Zhang M, et al. Genome-wide haplotype association study identify TNFRSF1A, CASP7, LRP1B, CDH1 and TG genes associated with Alzheimer’s disease in Caribbean Hispanic individuals. Oncotarget. 2015;6(40):42504–42514.

83. Ayers KL, Mirshahi UL, Wardeh AH, et al. A loss of function variant in CASP7 protects against Alzheimer’s disease in homozygous APOE ε4 allele carriers. BMC Genomics. 2016;17 Suppl 2:445.

84. Zhang X, Zhu C, Beecham G, et al. A rare missense variant of CASP7 is associated with familial late-onset Alzheimer’s disease. Alzheimers Dement. 2019;15(3):441–452.

85. Joutel A, Vahedi K, Corpechot C, et al. Strong clustering and stereotyped nature of Notch3 mutations in CADASIL patients. Lancet. 1997;350(9090):1511–1515.

86. Bianchi S, Zicari E, Carluccio A, et al. CADASIL in central Italy: a retrospective clinical and genetic study in 229 patients. J Neurol. 2015;262(1):134–141.

87. Abramycheva N, Stepanova M, Kalashnikova L, et al. New mutations in the Notch3 gene in patients with cerebral autosomal dominant arteriopathy with subcortical infarcts and leucoencephalopathy (CADASIL). J Neurol Sci. 2015;349(1-2):196–201.

88. Guerreiro RJ, Lohmann E, Kinsella E, et al. Exome sequencing reveals an unexpected genetic cause of disease: NOTCH3 mutation in a Turkish family with Alzheimer’s disease. Neurobiol Aging. 2012;33(5):1008.e17-e23.

89. Liu X, Zuo Y, Sun W, et al. The genetic spectrum and the evaluation of CADASIL screening scale in Chinese patients with NOTCH3 mutations. J Neurol Sci. 2015;354(1-2):63–69.

90. Patel D, Mez J, Vardarajan BN, et al. Association of Rare Coding Mutations With Alzheimer Disease and Other Dementias Among Adults of European Ancestry. JAMA Netw Open. 2019;2(3):e191350.

91. Wang G, Zhang D-F, Jiang H-Y, et al. Mutation and association analyses of dementia-causal genes in Han Chinese patients with early-onset and familial Alzheimer’s disease. J Psychiatr Res. 2019;113:141–147.

92. Sherman RM, Forman J, Antonescu V, et al. Assembly of a pan-genome from deep sequencing of 910 humans of African descent. Nat Genet. 2019;51(1):30–35.

93. Mills MC, Rahal C. A scientometric review of genome-wide association studies. Commun Biol. 2019;2:9.

94. Fereshtehnejad S-M, Vosoughi K, Heydarpour P, et al. Burden of neurodegenerative diseases in the Eastern Mediterranean Region, 1990-2016: findings from the Global Burden of Disease Study 2016. Eur J Neurol. 2019;26(10):1252–1265.

95. Babulal GM, Quiroz YT, Albensi BC, et al. Perspectives on ethnic and racial disparities in Alzheimer’s disease and related dementias: Update and areas of immediate need. Alzheimers Dement. 2019;15(2):292–312.

96. Marioni RE, Harris SE, Zhang Q, et al. GWAS on family history of Alzheimer’s disease. Transl Psychiatry. 2018;8(1):99.

97. Scott EM, Halees A, Itan Y, et al. Characterization of Greater Middle Eastern genetic variation for enhanced disease gene discovery. Nat Genet. 2016;48(9):1071–1076.

98. GenomeAsia100K Consortium. The GenomeAsia 100K Project enables genetic discoveries across Asia. Nature. 2019;576(7785):106–111.

99. Gao Y, Zhang C, Yuan L, et al. PGG.Han: the Han Chinese genome database and analysis platform. Nucleic Acids Res. 2020;48(D1):D971–D976.

100. Higasa K, Miyake N, Yoshimura J, et al. Human genetic variation database, a reference database of genetic variations in the Japanese population. J Hum Genet. 2016;61(6):547–553.

101. Genome of the Netherlands Consortium. Whole-genome sequence variation, population structure and demographic history of the Dutch population. Nat Genet. 2014;46(8):818–825.

102. Locke AE, Steinberg KM, Chiang CWK, et al. Exome sequencing of Finnish isolates enhances rare-variant association power. Nature. 2019;572(7769):323–328.

103. Martin AR, Kanai M, Kamatani Y, Okada Y, Neale BM, Daly MJ. Clinical use of current polygenic risk scores may exacerbate health disparities. Nat Genet. 2019;51(4):584–591.

