## Supplementary Material for "How understudied populations have contributed to our understanding of Alzheimer’s disease genetics"

### Supplementary data

#### Supplementary table 1. *APP, PSEN1* and *PSEN2* variants reported in each country with PubMed ID of study where available.

| Country | Gene | Unique variants | # Unique variants | Source | # Studies |
|---|---|---|---|---|---|
| Finland | *APP* | Ala673Thr | 1 | 31127772, 28556232, 23102935, 10854108, 10720282, 10643802, 9546792, 14759630, 11379823, 7550356, 9851443, 21959359 | 12 |
| | *PSEN1* | E9del, Glu318Gly, Met146Val, His163Arg | 4 | | |
| | *PSEN2* | | 0 | | |
| Argentina | *APP* | | 0 | 31153663, 9712537, 26166204 | 3 |
| | *PSEN1* | Thr119Ile, Met146Leu | 2 | | |
| | *PSEN2* | Asn141Ile | 1 | | |
| Australia | *APP* | Val717Ile, Leu723Pro | 2 | 26836186, 17632280, 9178856, 12192622, 9831473, 10665499 | 6 |
| | *PSEN1* | Ala79Val, E9del, Gln222His, Glu318Gly, Ile439Val/Ala713Thr, Leu219Pro, Leu271Val, Met146Ile, Met233Thr, Ser170Phe, Ser290Cys, Ser290Cys-E9del, His163Arg, Ser169Leu, Pro436Gln | 15 | | |
| | *PSEN2* | | 0 | | |
| Austria | *APP* | Val717Phe, Thr714Ile | 2 | 23415546, 15776278, 11487570 | 3 |
| | *PSEN1* | Ser170Phe | 1 | | |
| | *PSEN2* | | 0 | | |
| Belgium | *APP* | Glu682Lys, Ala713Thr, Lys724Asn, Ala692Gly | 4 | 11701593, 29859640, 21500352, 10548420, 16917905, 1303239 | 6 |
| | *PSEN1* | Leu282Val, Val198Leu, Ile143Thr, Gly384Ala | 4 | | |
| | *PSEN2* | Gly359Leufs*74 | 1 | | |
| Brazil | *APP* | | 0 | 28554858 | 1 |
| | *PSEN1* | Glu318Gly | 1 | | |
| | *PSEN2* | | 0 | | |
| Bulgaria | *APP* | | 0 | 19797784, Mehrabian et al., 2006 | 2 |
| | *PSEN1* | Leu381Val, Leu424Phe | 2 | | |
| | *PSEN2* | | 0 | | |
| Canada | *APP* | | 0 | 30138848, 27743520, 28034781, 27601731 | 4 |
| | *PSEN1* | Ala246Glu, Phe283Leu, Gly378Val | 3 | | |

| | | | | | |
|---|---|---|---|---|---|
| | *PSEN2* | | 0 | | |
| China | *APP* | Asp244Gly, Asp322Gly, Lys687Gln, Thr297Met, Val717Ile, Val695Met, Val715Met, Lys724Met, Met722Lys | 9 | 31385772, 31235344, 31235249, 30958370, 30822634, 30814350, 30440121, 29961914, 29304399, 29156377, 28269784, 28131463, 27926491, 27836335, 26422362, 25595498, 25323700, 24838186, 27816212, 30598257, 31440394, 15851849, 19853643, 24677022, 24737487, 25018108, 26402764, 26884997, 27838006, 30954774, 24650794, 12609057, 30090657 | 33 |
| | *PSEN1* | Ser169del, Ala246Glu, Arg157Ser, Arg269His, Arg352Cys, Gly111Val, Gly206Ser, Gly206Val, Gly378Glu, His214Arg, Ile167del, Ile249Leu, Leu173Trp, Leu226Phe, Leu262Phe, Leu286Val, Lys311Arg, Met139Ile, Met139Leu, Met233Leu, Met233Val, Phe105Leu, Phe105Val, Phe177Val, Phe386Ile, Phe388Leu, Pro433Ser, Thr147Ile, Tyr256Asn, Val103Gly, Val391Gly, His163Arg, Gln222Leu, Phe177Ser, Val97Leu, Phe105Cys, Leu262Ser, Gly394Val, Ala434Thr,Ala136Gly, Ile143Thr, Leu248Pro, Gly209Glu, Leu173Ser | 44 | | |
| | *PSEN2* | Arg163Cys, Asn141Tyr, Pro123Leu, Val139Met, His169Asn, Lys82Arg, Val214Leu, Val150Met, Asn141Asp, Ala379Asp | 10 | | |
| Colombia | *APP* | | 0 | 27372640, 30112632, 31026686, 30745123, 28550254, 26949549, 26619808, 24239249, 24239247, 22710270, 7550356, 9052708, 11568920, 12811988, 15230697, 18479822, 23134660, 23137948, 25471389, 20157243 | 20 |
| | *PSEN1* | Glu280Ala, Ile416Thr, Val94Met, Ile143Thr, Glu318Gly, Pro117Ala, Pro117Arg | 7 | | |
| | *PSEN2* | | 0 | | |
| Cuba | *APP* | | 0 | 12484344 | 1 |
| | *PSEN1* | Leu174Met | 1 | | |
| | *PSEN2* | | 0 | | |
| Caribbean-Hispanic | *APP* | Ser614Gly, Val340Met | 2 | 26214276, 25333068, 11710891, 23114514, 18797263, 27073747 | 6 |
| | *PSEN1* | Gly206Ala, Glu318Gly, Gly378Val | 3 | | |
| | *PSEN2* | Ile235Phe, Pro334Ala, Ala344Val | 3 | | |
| France | *APP* | Ala713Thr, Val717Ile, Lys724Asn, Asp694Asn, Glu693Lys, Ala692Gly, Val715Met, Ala235Val, Asp243Asn, Glu296Lys, Pro299Leu, Pro620Ala, Val715Ala, Leu723Pro | 14 | 28350801, 8863158, 28461250, 10874324, 8904759, 8634712, 9719376, 10200054, 11487570, 22475797, 16033913, 16941492, 27466472, 10441572, 26242991 | 15 |
| | *PSEN1* | Ala231Thr, Ala246Pro, Ala360Thr, Arg269His, Cys263Phe, E9-10del, Gln222His, Glu273Gly, Gly111Trp, Gly206Asp, Gly217Asp, Gly378Glu, Gly378Val, His163Arg, Ile143Thr, Ile180Asn, Leu153Val, Leu173Trp, Leu235Pro, Leu241Arg, Leu383Trp, Leu418Phe, Met139Lys, Met146Ile, Met210Arg, Met233Ile, Met233Thr, Met84Thr, Phe205_Gly206del;insCys, Phe237Cys, Phe237Leu, Pro117Gln, Pro264Leu, Pro88His, E9del, Tyr115Cys, Val391Phe, Ala79Val, Leu392Val, Cys410Tyr, Val82Leu, Tyr115His, Met139Thr, Leu262Val, Glu280Gly, Thr291Pro, Arg377Trp, Phe386Ser, Ser390Ile, Ser390Asn, Leu424His, Ile437Val, Glu69Asp, Thr116Ile, Pro117Ala, Glu120Asp, Ala260Val, Ala231Pro, Ser230Ile, Gln223Arg, His214Tyr, Ile213Thr, Glu184Gly, Phe177Leu, Trp165Cys, Met146Leu, Thr147Ile, Leu150Pro | 68 | | |
| | *PSEN2* | Thr122Pro, Arg284Gly, Ser130Leu, Lys161Arg, Met239Val | 5 | | |
| Germany | *APP* | Ile716Met, Val715Ala, Val717Ile, Val717Phe, | 6 | 26522186, 20457965, 20333730, 14648157, 10631141, 15776278, 15592140, | 21 |

| | | | | | |
|---|---|---|---|---|---|
| | | Val717Leu, Asp694Asn | | 9450781, 9521423, 9728730, 11487570, 15337637, 19073399, 29466804, 23850332, 7596406, 2025423, 26350633, 11409420, 23246540, 25130656 | |
| | *PSEN1* | Met146Leu, Ala79Val, Glu318Gly, Gly378Glu, Leu174Arg, Leu235Pro, Met139Val, Phe105Leu, Tyr115His, E9del, Gly209Val, His163Arg, Ser170Phe, Phe177Ser, Leu286Val, I238_K239insI, Phe176Leu | 17 | | |
| | *PSEN2* | Asn141Ile, Leu238Pro, Pro348Leu, Thr122Phe, Thr122Pro | 5 | | |
| Hungary | *APP* | Val717Phe | 1 | 28796010 | 1 |
| | *PSEN1* | | 0 | | |
| | *PSEN2* | | 0 | | |
| Iran | *APP* | Thr714Ala, Val717Ile | 2 | 29175279, 12034808, 22503161, 25138979 | 4 |
| | *PSEN1* | Val142Phe, Gly206Asp, His214Tyr | 3 | | |
| | *PSEN2* | | 0 | | |
| Iraq | *APP* | Ile716Phe | 1 | 31500908, 25182745 | 2 |
| | *PSEN1* | Val272Asp | 1 | | |
| | *PSEN2* | | 0 | | |
| Israel | *APP* | | 0 | 8931704 | 1 |
| | *PSEN1* | Glu120Asp | 1 | | |
| | *PSEN2* | | 0 | | |
| Italy | *APP* | Ala713Thr, Ala673Val, Val717Ile, Val715Met, Thr719Pro, Ile716Thr | 6 | 25174650, 9800154, 28532646, 10631141, 20164095, 26925509, 26549787, 25948718, 24718101, 23792692, 21422519, 20164579, 19286555, 18427071, 15365148, 15272895, 12037434, 7550356, 7611715, 7623584, 8513318, 7651536, 8805118, 10097173, 10822446, 11094128, 11126197, 14623725, 15006697, 15055444, 15755689, 16388371, 16902278, 16952411, 18525293, 19363265, 19584443, 20523046, 20842367, 21822699, 22531416, 27601731, 31177233, Terreni et al., 2002, Terreni et al., 2000, 15622541 | 46 |
| | *PSEN1* | Arg220Pro, His214Asn, Ile143Val, Ile408Thr, Met146Leu, Met84Val, Phe175Ser, Thr116Ile, Thr147Pro, Met233Thr, Leu392Val, Leu392Pro, Cys92Ser, Leu174Met, Leu166His, Glu318Gly, Gly394Val, Arg377Trp, Pro355Ser, Leu219Phe | 20 | | |
| | *PSEN2* | Met239Ile, Ser175Cys, Ala85Val, Met239Val, Ser130Leu, Thr122Arg | 6 | | |
| Japan | *APP* | Val717Ile, Asp678Asn, Val717Leu, Glu693del | 4 | 22882713, 18587238, 15732120, 15534188, 12111359, 11920851, 8737975, 29571857, 15364419, 12410385, 10447269, 9923762, 9292884, 8247223, 8741133, 8733303, 8945747, 8947284, 9007097, 9065558, 9109915, 9452052, 9521423, 10404731, 10644793, 12399144, 12686406, 12755040, 15201367, 16469444, 17968601, 30755281, 22572737, 22702962, 9804121, 19430857, 12391599, 24559647, 18300294,25743013, 23638752, 11561050, Higuchi et al., 2000 | 43 |
| | *PSEN1* | Glu123Lys, Glu184Asp, Gly209Arg, Gly217Asp, Gly266Ser, Gly378Glu, Leu286Val, Leu381Val, Leu392Val, Leu418Trp, Leu85Pro, Thr440del, Tyr154Asn, Leu420Arg, Val96Phe, Ile213Thr, His163Arg, Gly384Ala, Glu280Ala, Ala260Val, Ala285Val, E9del, Asn405Ser, Ala431Val, Leu250Val, Thr245Pro, Ile143Thr, Gly417Ser, Pro264Leu, Arg269His, Glu273Ala, Leu282Phe, Pro284Leu, Phe386Leu, Thr99Ala, His131Arg, Phe237Ile, Leu219Arg, Trp165Gly | 39 | | |
| | *PSEN2* | Ala258Val, Thr421Met | 2 | | |
| South Korea | *APP* | Glu145Lys, Val225Ala, Thr297Met, Pro484Ser, Val669Leu, Val715Met | 6 | 31623876, 30180983, 28532645, 28008242, 27785004, 31557888, 31308793, 31217084, 20213228, 18437002 | 10 |
| | *PSEN1* | Leu226Phe, Leu232Pro, Thr116Ile, Thr119Ile, Ala285Val, Glu184Gly, Gly209Ala, Gly417Ala, His163Pro, Leu232Pro, Thr116Ile, Trp165Cys, | 23 | | |

| | | | | | |
|---|---|---|---|---|---|
| | | Ala285Ser, Tyr389His, Tyr389Ser, Tyr115Cys, Glu120Lys, Met139Ile, Met233Thr, Gly206Ser, Ser170Phe, Tyr159Cys, His163Arg | | | |
| | *PSEN2* | His169Asn, Val214Leu | 2 | | |
| Malaysia | *APP* | Val717Ile | 1 | 28762277, 31557888, 26396515 | 3 |
| | *PSEN1* | Val96Phe, Glu280Lys | 2 | | |
| | *PSEN2* | Val68Glyfs*59 | 1 | | |
| Mexico | *APP* | | 0 | 16897084, 16628450, 9833068, 16801675, 9225696, 14966176 | 6 |
| | *PSEN1* | Ala431Glu, Leu171Pro, Thr354Ile, Asn135Asp, Leu235Val | 5 | | |
| | *PSEN2* | | 0 | | |
| Morocco | *APP* | c.1767_1768insC, c.1764_1765insC, c.1886_1887insC, c.1968delT, c.1881_1882insG, c.1932+2T>- | 6 | 24704512, 24627227, 9781063 | 3 |
| | *PSEN1* | Gly378fs, Glu318Gly | 2 | | |
| | *PSEN2* | E126fs, K306fs | 2 | | |
| Netherlands | *APP* | LysMet670/671AsnLeu, Glu693Gly, Ala692Gly | 3 | 24305500, 30797548, 9754958, 19494431, 9384602, 15258222 | 6 |
| | *PSEN1* | Leu418Phe, E9del, Ala79Val, Pro264Leu, His21Profs*2, Leu424Arg, Tyr115Cys, Ala231Val, Glu318Gly | 9 | | |
| | *PSEN2* | Met174Ile, Ala415Ser, Thr301Met | 3 | | |
| Peru | *APP* | | 0 | 24495933 | 1 |
| | *PSEN1* | Leu153Val | 1 | | |
| | *PSEN2* | | 0 | | |
| Poland | *APP* | Thr714Ala, Val715Ala | 2 | 14769392, 10337065, 9737546, 9507958, 12817569, 15003276, 15119739, 16546171, Gołąb-Janowska et al., 2009 | 9 |
| | *PSEN1* | Pro117Arg, Met139Val, His163Arg, Ile213Phe, Glu318Gly, Leu424Arg, Pro117Leu, Ala246Glu, Pro267Leu, Leu226Phe, Ile213Leu | 11 | | |
| | *PSEN2* | Gln228Leu | 1 | | |
| Portugal | *APP* | | 0 | 23489366, 18667258 | 2 |
| | *PSEN1* | Met164Val, Thr116Asn, Met233Thr, Ala260Val, Val272Ala | 5 | | |
| | *PSEN2* | | 0 | | |
| Spain | *APP* | Ile716Phe, Ala713Thr | 2 | 31204041, 26923015, 25638532, 23579331, 22906081, 22426017, 21501661, 21212633, 20858974, 20158511, 19001354, 18028191, 12925374, 11796781, 10768621, 22307680, 18667258, 21163230, 18637955, 27128372, 11723295, 9502232, 10025789, 10533070, 10732806, 11165779, 20157243, 12433263, 15488330, 18957849, 30838239 | 31 |
| | *PSEN1* | Arg220Gly, Glu120Gly, Glu318Gly, Gly209Glu, His214Asp, Ile439Ser, Leu166Arg, Leu173Phe, Leu235Arg, Leu248Arg, Leu282Arg, Leu286Phe, Leu286Pro, Leu392Val, Leu424Val, Lys239Asn, Met139Thr, Phe264Leu, Ser169Pro, Ser365Ala, Val261Leu, Val89Leu, ThrPro116/117SerThr, | 29 | | |

| | | | | | |
|---|---|---|---|---|---|
| | | Met233Leu, Ala409Thr, Phe105Val, His163Arg, Leu226Phe, Val272Ala | | | |
| | *PSEN2* | Met174Val, Thr430Met, Gly212Val, Asp439Ala, Val148Ile | 5 | | |
| Saudi Arabia | *APP* | Glu380Lys | 1 | 30636737, 22473143 | 2 |
| | *PSEN1* | Tyr195Cys, Arg377Thr | 2 | | |
| | *PSEN2* | Val139Met | 1 | | |
| Serbia | *APP* | Leu723Pro | 1 | 22221884 | 1 |
| | *PSEN1* | Gly206Asp, Arg108Gln | 2 | | |
| | *PSEN2* | Pro69Ala | 1 | | |
| Czech Republic | *APP* | | 0 | 28323683 | 1 |
| | *PSEN1* | Glu184Asp | 1 | | |
| | *PSEN2* | | 0 | | |
| South Africa | *APP* | | 0 | 14570818 | 1 |
| | *PSEN1* | Ile143Met | 1 | | |
| | *PSEN2* | | 0 | | |
| Sweden | *APP* | Glu693Gly, LysMet670/671AsnLeu | 2 | 29747683, 31386938, 20628413, 11528419, 1302033, 18413473, 9347932, 7550356, 7957938, 8117412, 8510829, 8619905, 8817335, 9605727, 19796846, 26836192, 9544835 | 17 |
| | *PSEN1* | Met146Val, His163Tyr, Ile143Thr, Glu318Gly, Thr116Asn, Leu262Phe, Met146Ile | 7 | | |
| | *PSEN2* | | 0 | | |
| Thailand | *APP* | Val717Ile, Val604Met | 2 | 10631141, 31557888, 30917570, 30510423 | 4 |
| | *PSEN1* | Glu184Gly | 1 | | |
| | *PSEN2* | | 0 | | |
| Tunisia | *APP* | | 0 | 26695639, 26145164 | 2 |
| | *PSEN1* | Ile83Thr | 1 | | |
| | *PSEN2* | | 0 | | |
| Turkey | *APP* | | 0 | 31296348, 22503161, 19912322 | 3 |
| | *PSEN1* | Leu424Pro, His163Arg, Pro264Leu, Leu134Arg, Leu262Val, Ala396Thr, Gln223Arg | 7 | | |
| | *PSEN2* | Met174Val, Ser130Leu | 2 | | |
| United Kingdom | *APP* | Val717Ile, His677Arg, Val717Leu, Lys496Gln, Ala713Thr, Val717Gly | 6 | 8247223, 25104557, 24880964, 12552037, 10090481, 7550356, 8742474, 9126060, 8733749, 9443865, 9521423, 10208579, 11084029, 16505331, 19667325, 21144619, 27249223, 23380992, 27777022, 30279455, 15115757, 26803359, 10401002, 24880964, 11405810 | 25 |
| | *PSEN1* | Ile168Thr, Leu166Val, Ser230Arg, Ile143Phe, Pro436Ser, Glu280Gly, Pro267Ser, | 46 | | |

| | | | | | |
|---|---|---|---|---|---|
| | | Met139Val,E9del, Leu250Ser, Glu120Lys, L113_I114insT, Glu120Asp, Ala426Pro, Pro264Leu, Leu219Pro, I83_M84del, Gly217Arg, Ile202Phe, Met146Ile, Tyr154Cys, Leu171Pro, Glu184Asp, Ile229Phe, Leu235Val, Phe237Leu, Ala260Val, Cys263Phe, Arg269His, Phe283Leu, Arg377Met, Gly378Val, Val393Phe, Cys410Tyr, Pro433Ser, Ala434Thr, Pro436Gln, L171_ L172insY (Leu171Tyr), Asn39Tyr, Arg42Leu, Tyr115Cys, Ser132Ala, Val142Ile, Ile227Val, Ile167del, Leu153Val | | | |
| | *PSEN2* | Ala237Val, Ser130Leu, Asp439Ala | 3 | | |
| United States | *APP* | Val717Ile, Ile716Val, Val717Phe, Asn660Tyr, Glu693Gly, Gly708Gly, Val642Phe, Asp694Asn | 8 | 29091718, 18413473, 21062519, 15159497, 15004326, 8247223, 20145736, 22312439, 1415269, 1611485, 7686976, 8154870, 9189043, 10963361, 11487570, 12112163, 16344340, 17366635, 27249223, 20375137, 31020001, 23752245, 11099448, 25812849, 7596406 | 25 |
| | *PSEN1* | Ala409Thr, Ala79Val, Arg269Gly, Arg352His, Gly206Ala, His214Tyr, Leu226Arg, Leu85Pro, Pro117Ser, Pro242His, Val412Ile, Y156F; Y156_R157insIY, Ala431Glu, Arg269His, Gly206Val, Ser170Phe, Met84Val, L113_I114insT, Thr147Ile, His163Arg | 20 | | |
| | *PSEN2* | Ala85Val, Asn141Ile, Met174Val, Leu238Pro, Lys115Glufs* | 5 | | |
| Denmark | *APP* | Ala673Thr, Thr714Ala | 2 | 26239177, 19659892, 9007311, 10439444, 18727676 | 5 |
| | *PSEN1* | Glu120Lys, Met146Ile, Thr116Asn | 3 | | |
| | *PSEN2* | Val393Met | 1 | | |
| Romania | *APP* | Val717Phe | 1 | 31282415, 9781063, 1925564 | 3 |
| | *PSEN1* | Leu85Pro, Glu318Gly | 2 | | |
| | *PSEN2* | | 0 | | |
| Uruguay | *APP* | | 0 | 23212405 | 1 |
| | *PSEN1* | Thr354Ile | 1 | | |
| | *PSEN2* | | 0 | | |
| Greece | *APP* | | 0 | 15776278, 15622541 | 2 |
| | *PSEN1* | Leu113Gln, Asn135Ser, Met146Leu | 3 | | |
| | *PSEN2* | | 0 | | |
| North American Aboriginal Kindred | *APP* | | 0 | 20481270 | 1 |
| | *PSEN1* | Leu250Phe | 1 | | |
| | *PSEN2* | | 0 | | |
| Ashkenazi Jewish | *APP* | | 0 | 21062519, 9781063 | 2 |
| | *PSEN1* | Cys410Tyr, Glu120Lys | 2 | | |
| | *PSEN2* | | 0 | | |

| | | | | | |
|---|---|---|---|---|---|
| African American | *APP* | Thr714Ile | 1 | 24413619, 12810495, 28106563, 15668448, 26888304, 17186461 | 6 |
| | *PSEN1* | Ile238Met, Met139Val, Ser167Phe, Pro267Ala, Asp333Gly | 5 | | |
| | *PSEN2* | Phe111Leu | 1 | | |
| Slovenia | *APP* | | 0 | 29930232 | 1 |
| | *PSEN1* | Pro264Ser, Phe105Cys | 2 | | |
| | *PSEN2* | | 0 | | |
| Ireland | *APP* | | 0 | 12370477 | 1 |
| | *PSEN1* | Ala431Val, Glu280Gly | 2 | | |
| | *PSEN2* | | 0 | | |
| Slovakia | *APP* | | 0 | 29404783 | 1 |
| | *PSEN1* | Thr116Asn | 1 | | |
| | *PSEN2* | | 0 | | |
| Taiwan | *APP* | Asp678His | 1 | | |
| | *PSEN1* | | 0 | | |
| | *PSEN2* | | 0 | 22558227 | 1 |
| India | *APP* | | 0 | | |
| | *PSEN1* | Trp165Cys | 1 | | |
| | *PSEN2* | | 0 | 29329714 | 1 |

**Supplementary table 2. Subset of supplementary table 1 to show *APP, PSEN1* and *PSEN2* variants unique to China to date.**

| Gene | Variants unique to China | Variants also reported in other countries |
|---|---|---|
| *APP* | Asp244Gly, Asp322Gly, Lys687Gln, Val695Met, Lys724Met, Met722Lys | Thr297Met, Val717Ile, Val715Met |
| *PSEN1* | Ser169del, Arg157Ser, Arg352Cys, Gly111Val, His214Arg, Ile249Leu, Lys311Arg, Met139Leu, Met233Val, Phe177Val, Phe386Ile, Phe388Leu, Tyr256Asn, Val103Gly, Val391Gly, Gln222Leu, Val97Leu, Leu262Ser, Ala136Gly, Leu248Pro, Leu173Ser | Ala246Glu, Arg269His, Gly206Ser, Gly206Val, Gly378Glu, Ile167del, Leu173Trp, Leu226Phe, Leu262Phe, Leu286Val, Met139Ile, Met233Leu, Phe105Leu, Phe105Val, Pro433Ser, Thr147Ile, His163Arg, Phe177Ser, Phe105Cys, Gly394Val, Ala434Thr, Ile143Thr, Gly209Glu |

| | | |
|---|---|---|
| *PSEN2* | Arg163Cys, Asn141Tyr, Pro123Leu, Lys82Arg, Val150Met, Asn141Asp, Ala379Asp | Val139Met, His169Asn, Val214Leu |

**Supplementary table 3. AD GWAS used to make pie chart.**

| PMID | Stage | Broad Ancestry |
|---|---|---|
| 17474819 | initial | European |
| 17553421 | initial | NOT REPORTED |
| 17553421 | replication | NOT REPORTED |
| 17975299 | initial | NOT REPORTED |
| 17998437 | initial | European |
| 17998437 | replication | European |
| 18449908 | initial | European |
| 18449908 | initial | NOT REPORTED |
| 18449908 | replication | European |
| 18823527 | initial | European |
| 18823527 | replication | NOT REPORTED |
| 18976728 | initial | European |
| 18976728 | replication | European |
| 19118814 | initial | European |
| 19118814 | replication | European |
| 19125160 | initial | European |
| 19136949 | initial | European |
| 19136949 | replication | European |
| 19734902 | initial | European |
| 19734902 | replication | European |
| 19734903 | initial | European |
| 19734903 | replication | European |

| | | |
|---|---|---|
| 20061627 | initial | European |
| 20452100 | initial | European |
| 20460622 | initial | European |
| 20460622 | initial | Hispanic or Latin American |
| 20460622 | replication | European |
| 20885792 | initial | Other |
| 20885792 | replication | NOT REPORTED |
| 21059989 | initial | African American or Afro-Caribbean |
| 21059989 | initial | European |
| 21059989 | initial | Hispanic or Latin American |
| 21059989 | replication | European |
| 21059989 | replication | Other |
| 21098978 | initial | Greater Middle Eastern (Middle Eastern, North African or Persian) |
| 21098978 | replication | European |
| 21116278 | initial | European |
| 21123754 | initial | European |
| 21379329 | initial | European |
| 21379329 | replication | European |
| 21379329 | replication | Hispanic or Latin American |
| 21390209 | initial | European |
| 21390209 | replication | NOT REPORTED |
| 21460840 | initial | European |
| 21460840 | replication | European |
| 21460841 | initial | European |
| 21460841 | replication | European |
| 21627779 | initial | European |

| | | |
|---|---|---|
| 21627779 | initial | NOT REPORTED |
| 21627779 | replication | European |
| 21627779 | replication | NOT REPORTED |
| 22005930 | initial | African American or Afro-Caribbean |
| 22005930 | initial | European |
| 22005930 | initial | Native American |
| 22005931 | initial | European |
| 22005931 | initial | NOT REPORTED |
| 22159054 | initial | African American or Afro-Caribbean |
| 22245343 | initial | European |
| 22430674 | initial | European |
| 22430674 | replication | European |
| 22785395 | initial | European |
| 22832961 | initial | European |
| 22832961 | replication | European |
| 22832961 | replication | NOT REPORTED |
| 22881374 | initial | European |
| 23150908 | initial | European |
| 23150908 | replication | European |
| 23150908 | replication | NOT REPORTED |
| 23374588 | initial | European |
| 23374588 | replication | European |
| 23419831 | initial | European |
| 23535033 | initial | European |
| 23562540 | initial | European |
| 23565137 | initial | East Asian |

| | | |
|---|---|---|
| 23565137 | replication | East Asian |
| 23565137 | replication | European |
| 23571587 | initial | African American or Afro-Caribbean |
| 23836404 | initial | NOT REPORTED |
| 23836404 | initial | European |
| 23836404 | replication | NOT REPORTED |
| 24162737 | initial | European |
| 24162737 | replication | European |
| 24755620 | initial | European |
| 24770881 | initial | NOT REPORTED |
| 24770881 | replication | NOT REPORTED |
| 24958192 | initial | European |
| 24958192 | replication | European |
| 24958192 | replication | NOT REPORTED |
| 25027320 | initial | European |
| 25027320 | replication | NOT REPORTED |
| 25043464 | initial | NOT REPORTED |
| 25043464 | replication | African American or Afro-Caribbean |
| 25043464 | replication | East Asian |
| 25043464 | replication | European |
| 25043464 | replication | NOT REPORTED |
| 25188341 | initial | NOT REPORTED |
| 25340798 | initial | NOT REPORTED |
| 25649651 | initial | NOT REPORTED |
| 25778476 | initial | NOT REPORTED |
| 25778476 | replication | European |

| | | |
|---|---|---|
| 26049409 | initial | East Asian |
| 26049409 | replication | East Asian |
| 26339675 | initial | Hispanic or Latin American |
| 26339675 | replication | Hispanic or Latin American |
| 26830138 | initial | European |
| 26913989 | initial | European |
| 26993346 | initial | NOT REPORTED |
| 27770636 | initial | African American or Afro-Caribbean |
| 28183528 | initial | African American or Afro-Caribbean |
| 28183528 | initial | East Asian |
| 28183528 | initial | European |
| 28183528 | initial | Greater Middle Eastern (Middle Eastern, North African or Persian) |
| 28183528 | replication | European |
| 28560309 | initial | European |
| 28714976 | initial | European |
| 28714976 | replication | European |
| 28870582 | initial | European |
| 29107063 | initial | African unspecified |
| 29107063 | initial | European |
| 29107063 | initial | Reported |
| 29274321 | initial | European |
| 29360470 | initial | European |
| 29360470 | replication | NOT REPORTED |
| 29615537 | initial | European |
| 29777097 | initial | European |
| 29777097 | initial | NOT REPORTED |

| | | |
|---|---|---|
| 29860282 | initial | European |
| 30201328 | initial | NOT REPORTED |
| 30201328 | replication | NOT REPORTED |
| 30413934 | initial | European |
| 30514930 | initial | European |
| 30617256 | initial | European |
| 30636644 | initial | European |
| 30805717 | initial | European |
| 30820047 | initial | European |
| 30820047 | replication | European |
| 31055733 | initial | European |
| 31473137 | initial | European |
| 31473137 | initial | NOT REPORTED |
| 31473137 | replication | European |
| 31473137 | replication | NOT REPORTED |

**Supplementary table 4. Counts of broad ancestries in AD GWAS used to make pie chart.**

| **Broad ancestry** | **# counts** |
|---|---|
| African American or Afro-Caribbean | 7 |
| African unspecified | 1 |
| East Asian | 6 |
| European | 85 |
| Greater Middle Eastern (Middle Eastern, North African or Persian) | 2 |
| Hispanic or Latin American | 5 |
| Native American | 1 |
| NOT REPORTED | 32 |
| Other | 2 |
